# Potentiation of drug toxicity through virus latency reversal promotes preferential elimination of HIV infected cells

**DOI:** 10.1101/2022.01.12.476003

**Authors:** Thanh Tung Truong, Manuel Hayn, Camilla Kaas Frich, Lucy Kate Ladefoged, Morten T. Jarlstad Olesen, Josefine H. Jakobsen, Cherie K. Lunabjerg, Birgit Schiøtt, Jan Münch, Alexander N. Zelikin

**Affiliations:** Dr. T.T. Truong, Dr. C. Kaas Frich, Dr. M.T. Jarlstad Olesen, C.K. Lunabjerg, Prof. B. Schiøtt, Prof. A.N. Zelikin Department of Chemistry, Aarhus University, Langelandsgade 140, Aarhus C 8000, Denmark; Institute of Molecular Virology, Ulm University Medical Center, 89081 Ulm, Germany; Department of Biomedicine, Aarhus University, Aarhus 8000, Denmark; iNano Interdisciplinary Nanoscience Centre, Aarhus University, Aarhus 8000, Denmark

**Author notes:** authors contributed equally (Thanh Tung Truong, Manuel Hayn).

**Keywords:** HIV latency, drug design, latency reversal

## Abstract

Eliminating latently infected cells is a highly challenging, indispensable step towards the overall cure for HIV/AIDS. We recognized that the unique HIV protease cut site (Phe-Pro) can be reconstructed using a potent toxin, monomethyl auristatin F (MMAF), which features Phe at its C-terminus. We hypothesized that this presents opportunities to design prodrugs that are specifically activated by the HIV protease. To investigate this, a series of MMAF derivatives was synthesized and evaluated in cell culture using latently HIV-infected cells. Cytotoxicity of compounds was enhanced upon latency reversal by up to 11-fold. In a mixed cell population, nanomolar concentrations of the lead compound depleted predominantly the HIV-infected cells and in doing so markedly enriched the pool with the uninfected cells. Despite expectation, mechanism of action of the synthesized toxins was not as HIV protease-specific prodrugs, but likely through the synergy of toxicities between the toxin and the reactivated virus.

---

Eradication of human immunodeficiency virus (HIV) from the human body is a grand, unmet medical need. HIV is the causative agent of the acquired immunodeficiency syndrome (AIDS) and currently affects over 37 million people worldwide (UNAIDS report 2018). Combined anti-retroviral therapy (ART) prevents HIV replication and the development of AIDS. Consequently, the life expectancy of HIV-infected individuals receiving ART is near-identical to that of uninfected individuals. However, ART has side effects, is expensive, and requires strict adherence to the drug regimen. In addition, patients have to take ART for their entire life because it fails to eradicate the virus from the body. The major reason for this is that HIV integrates into the genome of its host cells and forms latent viral reservoirs in long-lived resting or quiescent memory CD4 + T cells.^1–2^ ^3^ These cells carry stably integrated, transcriptionally silent, but replication-competent proviruses. While these cells do not produce viral particles in their resting state, they can give rise to infectious virions upon activation.^4–6^ If this occurs after ART is interrupted, HIV rebounds. So far, only two HIV infected individuals have been cured of HIV/AIDS, both after allogenic transplantation of hematopoietic stem cells from donors with a homozygous mutation in the HIV coreceptor CCR5. ^7–8^ However, this approach is highly risky and not broadly applicable. As it stands, a practical cure is far beyond reach. A successful cure of HIV would be a formidable achievement that might also pave the way to curing other persistent viral infections.

One strategy that is intensively investigated towards the HIV/AIDS cure is the “shock and kill” approach. Initially, treatment with so-called latency-reversing agents (LRAs) reactivates latent HIV hiding in immune cells (the “shock” step). Consequently, virus producing cells can be targeted and eliminated by the human immune system while new HIV infections are prevented by ART.^9–10^ However, the “kill” step relies on the human immune system and is insufficient to achieve efficient elimination of all HIV infected cells.^11^ ^12^ The fundamental challenge of HIV cure would therefore benefit tremendously from the development of therapeutic “kill” agents, to specifically eliminate the virus infected cells. Yet this endeavour has proven to be highly challenging and decades of research on this subject delivered only solitary examples of success.

A therapeutic strategy that proved to be highly promising focused on the development of “pathogen-activated antiretroviral treatment”,^13–17^ designed with the knowledge of the unique substrate specificity of the HIV protease. HIV protease-specific peptide sequences have been used to inactivate cytotoxic proteins (through the design of zymogens^15–17^ and protein drug carriers^14^) whereby activity of the protease liberated the therapeutic agents ensuing death of the cell that harbors HIV. The developed opportunities are fundamentally exciting but require intracellular delivery of protein therapeutics which *in vivo* remains a major challenge on its own. ^18–19^

In this work, we approached the design of the medicinal “kill” agent to eliminate the virus-infected cells from a different angle. Instead of considering the full peptide sequences specific to the HIV protease, we focused on the unique HIV protease cleavage site between the aromatic amino acids phenylalanine or tyrosine and proline (Phe, Tyr and Pro, respectively).^20^ We realized that a clinically validated toxin, monomethyl auristatin F (MMAF), contains a C-terminal phenylalanine, and lends itself to reconstruct the Phe-Pro cut site (Figure 1A). Pro derivatives of MMAF could therefore potentially serve as prodrugs if the potent toxin would be released by the activity of the HIV protease, thus ensuing cell killing of the virus-infected cells. MMAF has low cell membrane permeability and hence a low bystander cell killing effect,^21^ further contributing to the attractiveness of this drug. We synthesized a total of eight Pro-containing, C-substituted derivatives of MMAF (Figure 1A) whereby “R” was chosen from a range of functionalities to optimize the compound cell entry, and at the same time to optimize binding to the hydrophobic S2’ pocket of the protease (S1’ being occupied by proline). For synthesis details and compound characterization, see Supplementary Information.

**Figure 1:**
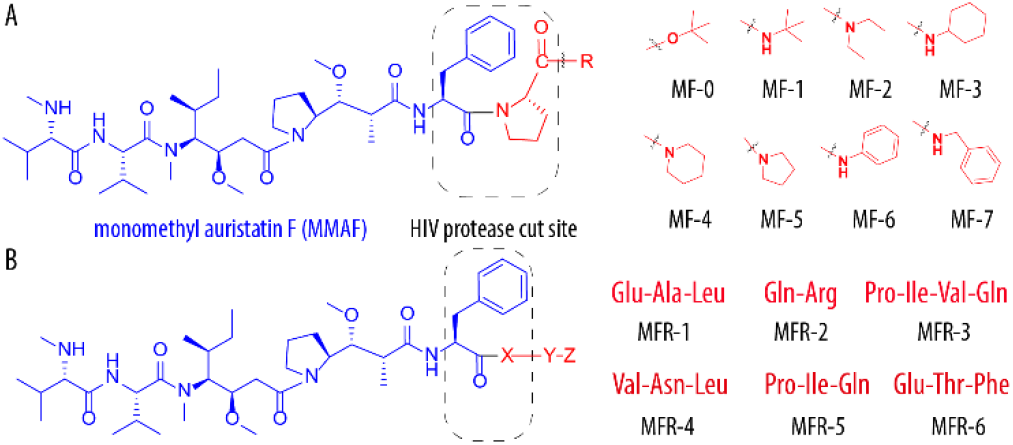
Chemical formulas of the derivatives of MMAF (A) engineered such as to reconstruct the Phe-Pro HIV protease cut site and to include the R group to optimize protease binding; and (B) engineered using MMAF toxin and peptidic motifs from the known HIV protease peptide substrates.

From a different perspective, we also considered that the HIV protease is a symmetrical homodimer with a bidirectional binding of the substrate,^20^ and considered that a range of commercially available HIV-specific peptide sequences feature Phe (or Tyr) in their structure at the cut site. We envisioned that, using the fragment-based drug design methodology, it is possible to preserve the sequence “right” of the cut site of the HIV-protease substrate, to maintain the affinity for the protease, and use MMAF as the “left” fragment, to afford an HIV-specific prodrug for the toxin (Figure 1B). To realize this synthetically, the second series of prodrugs (MFR series) was designed to contain the MMAF toxin connected at the C-terminus to EAL (derived from KVNLEAL, commercialized as an HIV protease-specific peptide by Sigma), QR (from the YIF-QR peptide from Bachem); and PIVQ (from SQNY-PIVQ, commercialized by Anaspec). In this design, we also included fragments of the three sequences that were identified in the in silico-based modelling of the peptides with HIV protease specificity,^22^ namely (F)ETF, (Y)VNL, and (F)PIQ. For synthesis details and compound characterization, see the Supplementary Information.

To study the newly synthesized derivatives of MMAF, we first established a robust cell culture model for latency reversal. Four latently infected Jurkat-based T cell lines containing a full-length integrated HIV-1 genome that expresses GFP upon activation (J Lat 8.4, 9.2, 10.6, and 11.1) ^23^ were analyzed for latency reversal using the histone deacetylase inhibitor vorinostat (SAHA) or tumor necrosis factor alpha (TNFα), an immune stimulatory cytokine. Latency reversal was quantified 48 h post-treatment using flow cytometry to monitor GFP expression. In parallel, we determined cell viability using a fluorescent viability dye. SAHA concentrations exceeding 1.25 μM caused more than 50% reduction in cell viability in all analysed J-Lat cell clones (Figure S1A). Concentrations of SAHA below or equal to 1.25 μM, however, induced no or only moderate levels of GFP expression (Figure S1B). In contrast, treatment with TNF-α was efficacious in terms of virus latency reversal using TNF-α concentration ≥5 ng/mL and devoid of noticeable toxicity at concentrations ≤10 ng/mL (Figure S1C,D). In all further experiments, latency reversal was achieved using 10 ng/mL cytokine concentration.

We determined the cytotoxic activity of MMAF and derivatives thereof designed in this work in J-Lat 10.6 cells. Synthesized reagents proved to be potent toxins with toxicity-related IC_50_ values as low as 0.5 nM (Figure 2, for numerical values see Table 1). For the Pro-based reagents (MF series), typical IC_50_ values were between 4 and 20 nM and those for the MFR series between 60 and 570 nM. Noteworthy, for the MF series, IC_50_ values were lower than that for MMAF, as is readily explained by the masking effect of the Pro-R group to the highly polar carboxylate group of MMAF.^21^ Nevertheless, these data also illustrate that the synthesized compounds/prodrugs do not fully prevent cell killing by the toxin. Altogether, this observation is not surprising and in agreement with prior results on MMAF derivatives.^21^

**Table 1.**
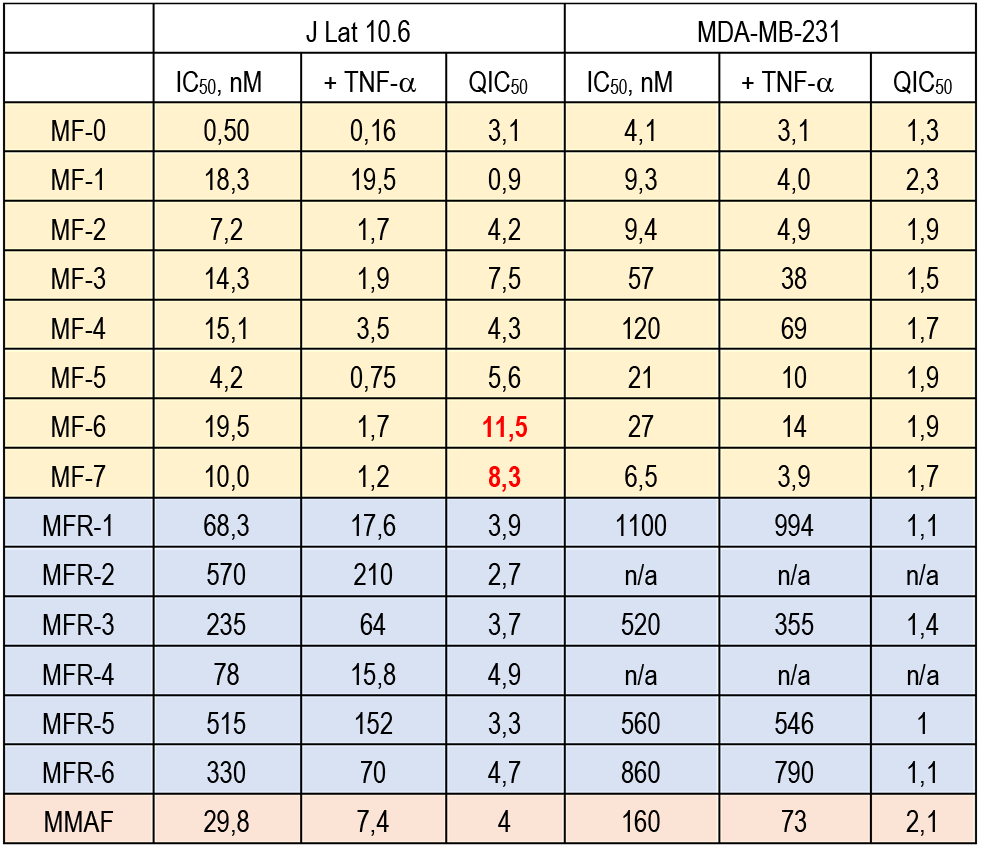
Toxicity related values of IC_50_ for MMAF and the developed compound library in J-Lat 10.6 cells harboring latent HIV and in MDA-MB-231 cancer cells (no latent HIV), in the presence or absence of HIV latency reversal agent, TNF-α. QIC_50_ is defined as a ratio of respective IC_50_ values for the same drug in cells without and with addition of TNF-α and reflects fold-potentiation of the drug toxicity upon addition of the HIV latency reversal agent. Presented values are based on at least 3 independent experiments and presented as mean *±* SD.

**Figure 2:**
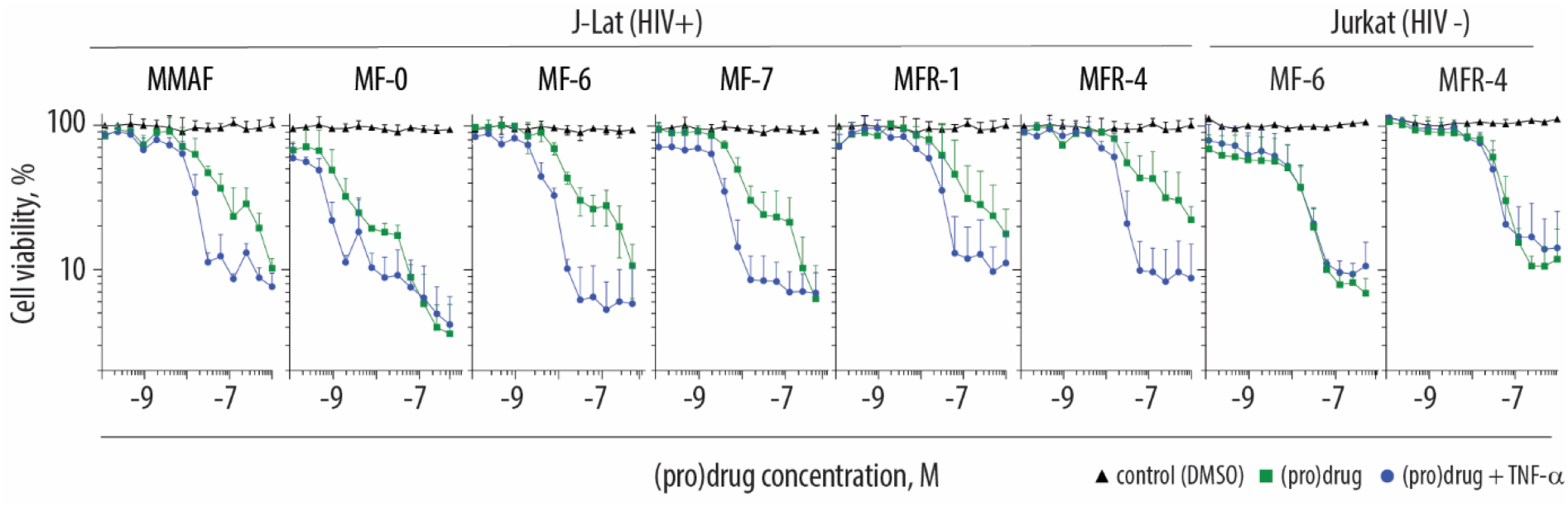
Cytotoxicity of MMAF and the developed compound library in the J-Lat 10.6 cells (latently infected with HIV) or the parent non infected Jurkat cells in the presence and absence of the latency reversal agent, TNFα. Shown are the mean values of three independent experiments ± SD.

Next, we quantified drug toxicity under conditions of HIV latency reversal. For this, J-Lat 10.6 cells were treated with TNF-α and subsequently exposed to the library compounds. In the presence of TNF-α, IC_50_ values for the parent MMAF declined, illustrating that the cytokine exhibits an unexpected, albeit minor, non-specific drug potentiation effect (Table 1). We define the value of QIC_50_ as the ratio of the corresponding IC_50_ values obtained for the toxin in the absence or the presence of TNF-α (Table 1); this ratio for MMAF in J-Lat 10.6 cells was 4. In these cells, the overall majority of the synthesized toxins was characterized by similar QIC_50_ values, suggesting that latency reversal leads to a minor potentiation of toxicity of treatment. For control, drug toxicity was also quantified using a breast cancer line MDA-MB-231, in which case the QIC_50_ values were also in the range between 1 and 2, revealing minor, non-specific effects mediated by TNF-α.

Two synthesized toxins, MF-6 and MF-7, revealed QIC_50_ values of 11.5 and 8.3 in J-Lat cells and only QIC_50_ of ~ 2 in the cancer cell line. Thus, in cancer cells, effects of TNF-α were similar to those observed for all the synthesized reagents. In contrast, in the J-Lat cells harbouring latent HIV genome, TNF-α treatment induced a significant potentiation of the compounds’ effect. A common design feature for the two lead compounds is that both MF-6 and MF-7 are MMAF derivatives extended with proline, not the di/tripeptide sequence; and that proline is further protected into an aromatic amide (whereas all other MF series agents were aliphatic amides). MF-6 and MF-7 thus share structural similarity with lopinavir, ritonavir and other protease inhibitors that contain aromatic binding groups (mimicking Phe) on either side of the hydroxyethylene spacer, that is, on either side of the symmetry plane when the drug binds to the symmetrical, bidirectional binding site of the HIV protease.^24^

To investigate the potential of specific killing of the HIV-infected cells, activity of MF-6 and MF-7 was analyzed via flow cytometry, distinguishing between the GFP+ cells with latency reversed virus and GFP-cells that are devoid of transcriptionally active HIV. Latency reversal in J-Lat 10.6 cells resulted in a mixed population of cells with an approximately 2:1 ratio of GFP+/GFP-cells (Figure 3A). At compound concentrations below 3-5 nM, we observed a strong cell-killing effect in the GFP+ (HIV+) cells, and a much less pronounced change in viability for the GFP-(HIV-) cells. Drug dosing afforded predominant elimination of the HIV+ cells and upon treatment, the cell pool is significantly enriched with the HIV-negative cells (Figure 3B). These data comprise an important step towards the development of a therapeutic “kill” agent for the “shock and kill” strategy.

**Figure 3.**
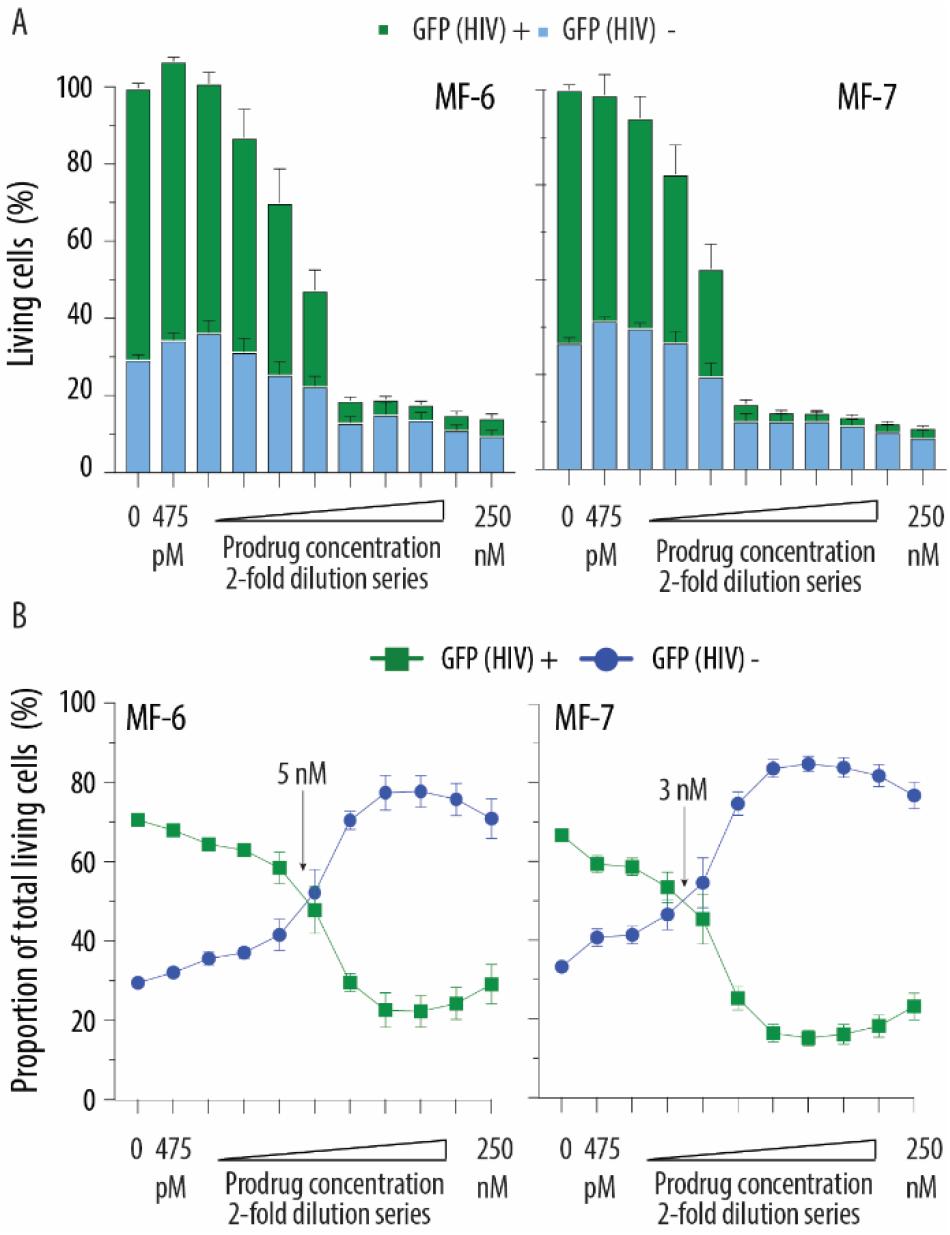
Differential killing potential for the MF6 and MF7 prodrugs in latently infected J-Lat cells in the presence of 10 μg/L TNF-α (latency reversal agent) as a function of prodrug concentration: (A) Stack-bar view whereby all viable cells are color-coded as GFP (HIV) positive or negative; (B) proportion of total live cells that test positive or negative for GFP (HIV). Shown are the mean values of three independent experiments each performed in triplicates ± SEM.

In an effort to validate the compound binding to the HIV protease, we used computational methods and performed induced fit docking calculations for MF-7 using atomic coordinates for the HIV protease co-crystallized with lopinavir. ^25^ The calculated binding mode of MF-7, unlike that of MMAF taken as a control, occurred such as to occupy the protease binding site, thus providing computational support of the experimental prodrug toxicity data. Indeed, while pristine MMAF exhibited superficial binding to the protease, docking calculations suggested that MF-7 binds such as to overlap significantly with lopinavir (Figure 4). This was true either with or without a structural water molecule included in the calculations. The calculated binding mode matched well with the expected compound fitting into the protease active site, specifically in that Pro is positioned in the immediate vicinity of Asp 25/25’ (the protease cut site), and the aromatic rings of the compound are largely co-localized with the two phenyl and isopropyl side chains of lopinavir (for details and additional figures, Supporting Information).

**Figure 4:**
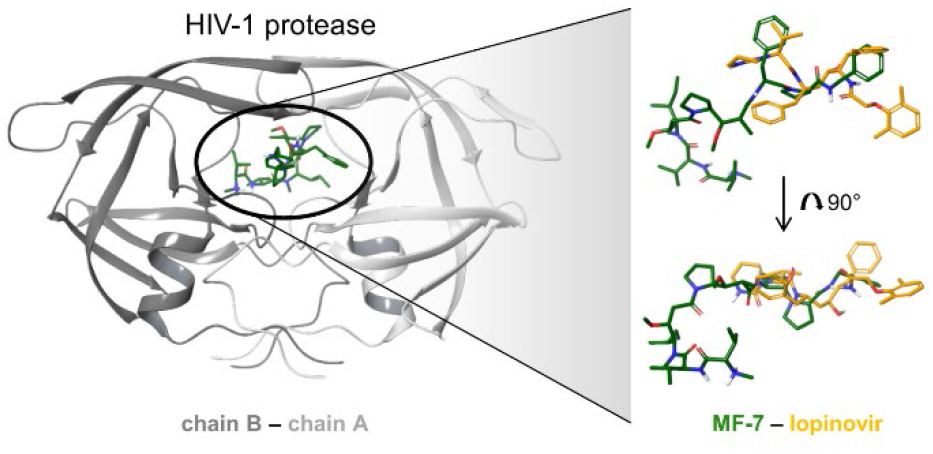
Computational illustration of MF-7 binding to HIV-1 protease. Shown here is the most favorable binding mode of MF-7 in HIV-1 protease (SP Gscore = −6.0 kcal/mol) when docked with a bridging water molecule. Structures to the right illustrate the protease-bound MF-7 and lopinavir (from PDB entry 6DJ1).

Next, we performed an evaluation of the compound library in the context of the protease-mediated drug release. We observed the expected activity of the HIV protease on the commercially available fluoregenic substrate via both, HPLC and the fluorescence based readout (Figure 5). However, despite expectations, we registered no release of MMAF from the compounds synthesized in this work in the presence of the HIV protease (data for MF-7 shown in Figure 5). In the protease inhibition experiments with a fluorescence read-out, we observed that upon addition of MF-7, conversion of the fluoregenic substrate was decreased, likely indicating competitive enzyme inhibition. This effect was statistically significant, although its magnitude was rather modest.

**Figure 5.**
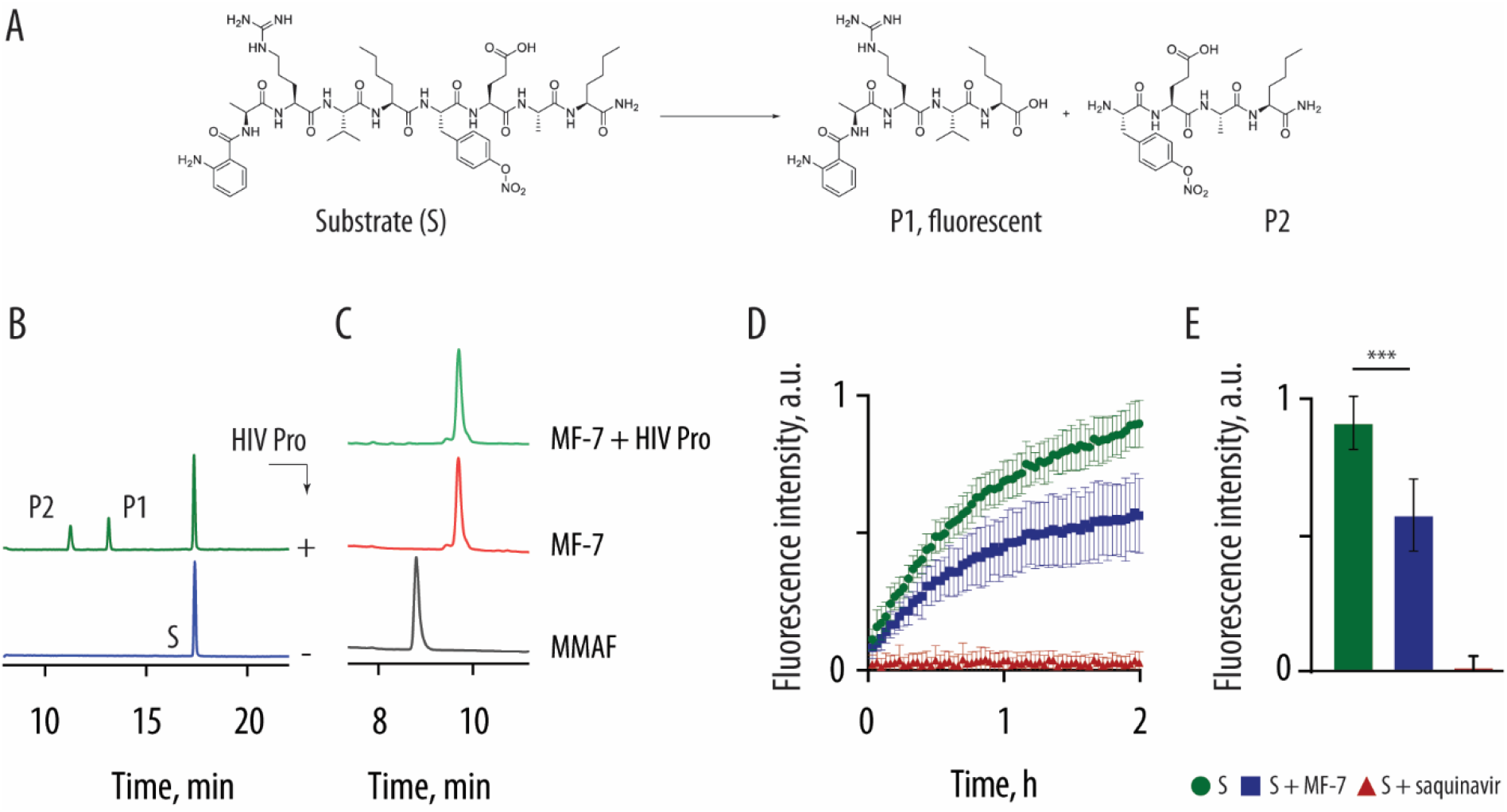
Evaluation of MF-7 as a substrate or an inhibitor for the HIV protease. (A) Chemical formula of the fluorogenic substrate to the HIV protease (Sigma product H0790) and its corresponding products of proteolytic cleavage; (B) HPLC elution profile illustrating the expected enzymatic scission of the substrate (incomplete conversion); (C) HPLC elution profiles for MMAF and MF-7, for the latter with and without a treatment of the HIV protease; (D,E) Fluorescence of solutions containing the HIV protease and its fluoregenic substrate in the presence of MF-7 or saquinavir, in kinetic mode (D) or end-point measurement (E). Panels D,E: shown are results based on three independent experiments; statistical evaluation was carried out via two-way ANOVA (GraphPad Prism v. 9), *p* < 0,001.

Taken together, computation methods, quantification of drug release, and the protease inhibition studies support the compound binding to the protease, albeit with modest affinity and without enzymatic activity on MF-7. Thus, most likely, the synthesized reagents exert their preferential toxicity upon HIV latency reversal through a mechanism different to the originally proposed activity of the prodrugs being activated by the HIV protease. This conclusion is further strengthened by the observation that potentiation of the toxin upon HIV latency reversal (Figure 2 and Table 1) was not suppressed by the HIV protease inhibitor (saquinavir).

One plausible mechanism of the observed enhanced drug toxicity in the HIV+ cells is based on a possible synergy of cytotoxic effects exerted by the synthetic toxin and the virus. Indeed, synergy between viral cytopathic effects (CPE) and drug-induced toxicity is well-established as a means of the combination cancer therapy and is investigated in clinical trials.^26–30^ Furthermore, chemotherapy clinical trials in the HIV+ patients also documented a significant decrease of the viral load in blood, indicative of elimination of the virus-producing cells.^31^ Synergy between chemotherapy and CPE therefore merits investigation as an approach to eliminate the virus-infected cells. Specifically, more work is required to enhance the QIC_50_ to highest possible values, and to identify the exact mechanism of enhanced toxicity in the HIV+ cells.

## EXPERIMENTAL SECTION

For details on compound synthesis and characterization, see Supplementary Information.

### Cell culture

J-Lat 11.1, 10.6, 9.2 and 8.4 were obtained from the NIH AIDS Reagent Program and maintained in RPMI1640 medium supplemented with FCS (10% (v/v)), L-glutamine (2 mM), streptomycin (100 mg/mL) and penicillin (100 U/mL). Cells were cultured at 37°C, 90% humidity and 5% CO2. The cells were split 1:20 or 1:30 regularly twice a week. Parental Jurkat cells were obtained from ATTC (Clone E6-1; ATCC® TIB-152™). Cell were cultured in RPMI1640 medium supplemented with FCS (10% (v/v)), L-glutamine (2 mM), streptomycin (100 mg/mL) and penicillin (100 U/mL). Cells were cultured at 37°C, 90% humidity and 5% CO2. The cells were split 1:10 or 1:20 regularly twice a week.

### Toxicity treatment

75,000 J-Lat or parental Jurkat cells were incubated in 96 U-Well microtiter plates with the respective compound dilutions prepared in RPMI medium in a final volume of 200 μL per well. Cells were then either stimulated with 10 ng/mL TNFα or not in the absence or presence of MMAF and prodrugs. Cells were incubated at 37°C, 5% CO2 and 90% humidity for 2 days. 48 hours post treatment, cell viability was determined as described below.

### Flow cytometry analysis of cell viability

Cells were spinned down at 350 × g and room temperature for 3 minutes. Supernatants were discarded. Cells were washed twice in 1x PBS. eBioscience™ Fixable Viability Dye eFluor™ 780 (Thermo Scientific, #65-0865-14) was diluted 1:1000 (v/v) in 1x PBS. Cells pellets were resuspended in 50 μL in the solution containing the viability dye and incubated at room temperature for 15 minutes in the dark. Cell were washed twice in 1x PBS. Finally, cells were fixed in 4% PFA at 4°C for 1 hour. J-Lat and parental Jurkat cells were gated based on forward and side scatter characteristics, followed by exclusion of doublets and then by the viability dye positive and negative cells. Data were generated with BD FACS Diva 6.1.3 Software using the FACS Canto II flow cytometer. Data analysis was performed using FlowJo 10.6 Software (Treestar).

### Cell Titer Glo viability assay

Cells were spinned down at 350 × g and room temperature for 3 minutes. Supernatants were discarded. Cells were washed twice in 1x PBS. The CellTiter-Glo® Luminescent Cell Viability Assay Kit (Promega, # G7570) was used as recommended in the manufacturer’s protocol. Cell viability was quantified as relative light units (RLU) per second with an Orion Microplate luminometer (Berthold).

### Computational Methods

#### Protein preparation

The HIV-1 protease structure was obtained from PDB entry 6DJ1 (Ref^25^) and prepared for docking calculations using the Protein Preparation Wizard ^25, 32^ (available in Maestro, Schrödinger Suite 2019, Schrödinger, LLC, New York, NY, 2019). This template was selected due to it having been co-crystallized with lopinavir, which contains a PheProPhe isostere similar to MF-6 and MF-7, and the conformation of the flap domains was therefore expected to be compatible with MF-6 and MF-7 binding. All ions and water molecules were deleted except for one water molecule bridging the co-crystallazed ligand lopinavir and Ile50. When dual conformations were present in the crystal structure, the most populated conformation of residues and ligand was selected. The protonation states of titratable residues were evaluated using PROPKA ^33^ at neutral pH resulting in Asp25’ (chain A) being modelled in its protonated state, which is in accordance with neutron scattering data. ^34^ All other residues were modelled in their default states. The protein was then subjected to a structure minimization restrained to a maximum heavy atom RMSD of 0.3 Å.

#### Ligand preparation

The structure of MMAF was extracted from PDB entry 5J2U. ^35^ Atom types and bond orders were assigned manually and the stereochemistry was checked in Maestro (Schrödinger Suite 2019, Schrödinger, LLC, New York, NY, 2019). The protonation state of relevant functional groups was assessed ^36^ using Epik resulting in both termini being modelled as charged. The structure was then minimized and submitted to a conformational search using MacroModel available within Maestro, the OPLS 2005 force field ^37^ and otherwise default settings. The lowest energy conformation was applied in the docking calculation and as starting point for generating the chemical structures of MF-6 and MF-7. The MMAF structure was manually extended to MF-6 and MF-7, respectively, using the build panel available in Maestro, and prepared analogously to MMAF as described above.

#### Docking calculations

Docking calculations were performed using the *extended* induced fit docking protocol ^38^ which utilizes three steps to achieve both ligand and protein flexibility (Schrödinger Suite 2019, Schrödinger, LLC, New York, NY, 2019). In the first step, the ligand is docked using a soft potential to allow for an initial binding mode despite a poor protein/ligand fit. In this step, the binding site is initially evaluated and residues protruding into the binding site or highly flexible residues are temporarily mutated to alanine as to not obstruct initial binding. The *extended* protocol includes automated mutation of problematic residues and performs docking calculations on each resulting protease mutant. In the second step, any mutated residues are changed back to the original amino acid and all side chains in the binding site (5Å around docked ligand) are optimized to accommodate the docked ligand. In step three, the docking step is repeated using the optimized binding site and the full potential function. The binding site center was defined as the centroid of lopinavir and the binding site size was set to 46 Å. The first docking step was allowed to return a maximum of 80 poses, while the second docking step could return a maximum of 20 poses. Both docking steps were performed using the standard precision scoring function. ^39^ The docking calculation was repeated without the structural water molecule near Ile50.

### Studies with the HIV Protease

To confirm activity of the HIV protease a positive control was run alongside all drug-release experiments using an HIV Protease fluoregenic substrate (Sigma Aldrich Cat. No H0790).

For HPLC quantification of drug release, stock solutions of the compounds in DMSO (10 g/L) were diluted to a final concentration of 200 μM in acetate buffer (100 mM NaOAc, 1 M NaCl, 1 mM EDTA, 1 mM DTT, pH 4.7) and incubated at 37 °C for 24 h in the presence or absence of HIV-1 protease (83 nM). 20 μL of each reaction mixture was added to MeOH (80 μL) to precipitate the HIV protease. The MeOH layer was analyzed via HPLC. All release studies were conducted in triplicates.

The protease inhibition studies were carried out in 100 mM MES buffer (pH 5,5) supplemented with 400 mM NaCl, 1 mM EDTA, 1 mM DTT, and 1 g/L BSA. Reagent concentrations were 80 nM HIV-PR, 1 μM HIV protease fluorescent substrate, 10 μM MF-7, 1 μM Saquinavir. Experiments were conducted at 37 C with shaking, reading solution fluorescence every 2 min with excitation and emission wavelengths of 320 and 420 nm, respectively.

## Acknowledgements

This work was supported by amfAR, The Foundation for AIDS Research (to ANZ, grant No 109328-59-RGRL), the Lundbeck Foundation (to ANZ, grant No R108-A10354), the Danish Council for Independent Research, Technology and Production Sciences, Denmark (to ANZ, grant No DFF −4184-00177), and CRC 1279 (to JM). We thank Drs. S. Harris Wibowo, M. Tolstrup, P. Denton and A.H.F. Andersen for involvement in the early stages of the project.

## TOC image

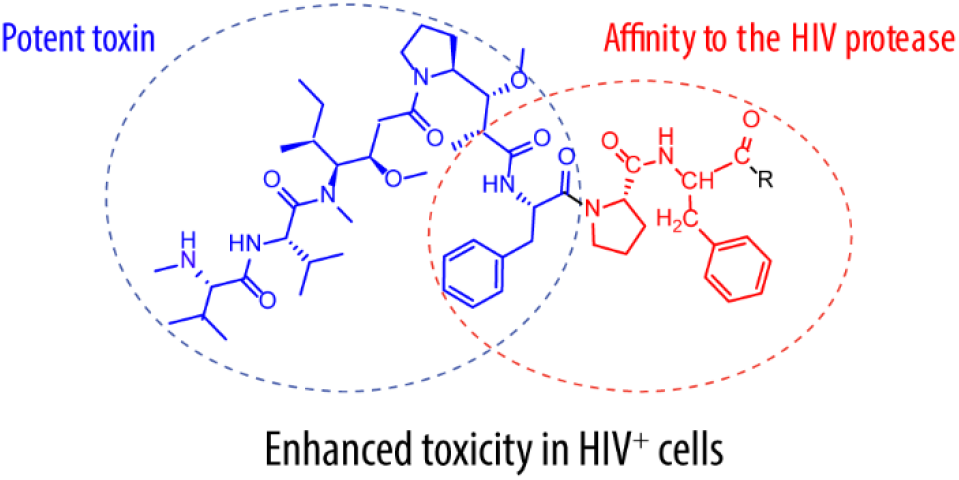

## Supporting Information

### SUPPLEMENTARY FIGURES

**Figure S1:**
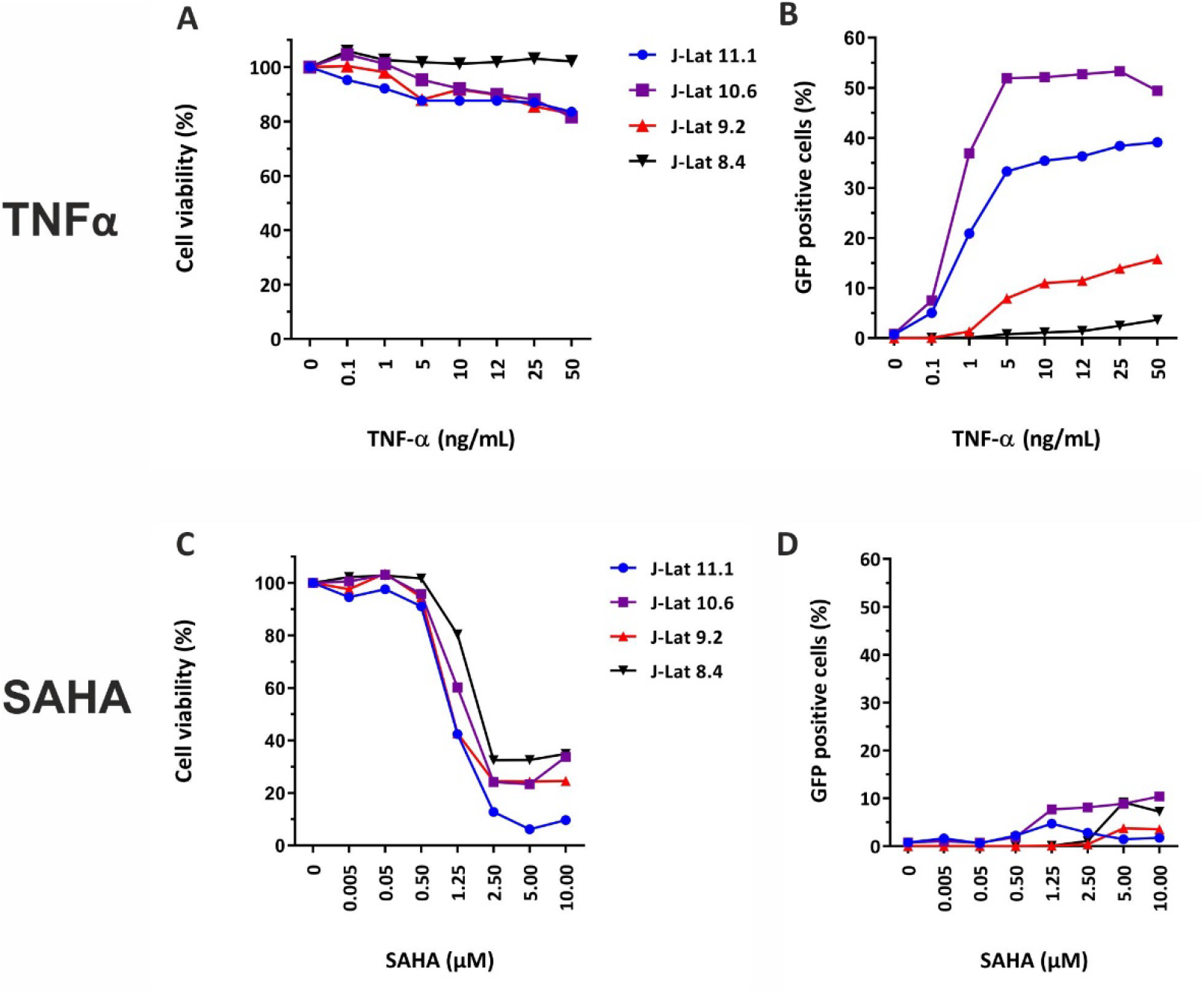
Reactivation of latent HIV-1 in J-Lat cells in presence of TNFα or Vorinostat (SAHA). J-lat cells were inucbated with TNFα or SAHA at the indicated concentrations for 48 hours. (A) and (C): Cell viability was determined by additionally staining all cells with the eFluor780 fixable viability dye according to the manufacturer‘s recommendations. (B) and (D): Reactivation of latent HIV-1 was measured by analyzing GFP expression via flow cytometry.

**Figure S2.**
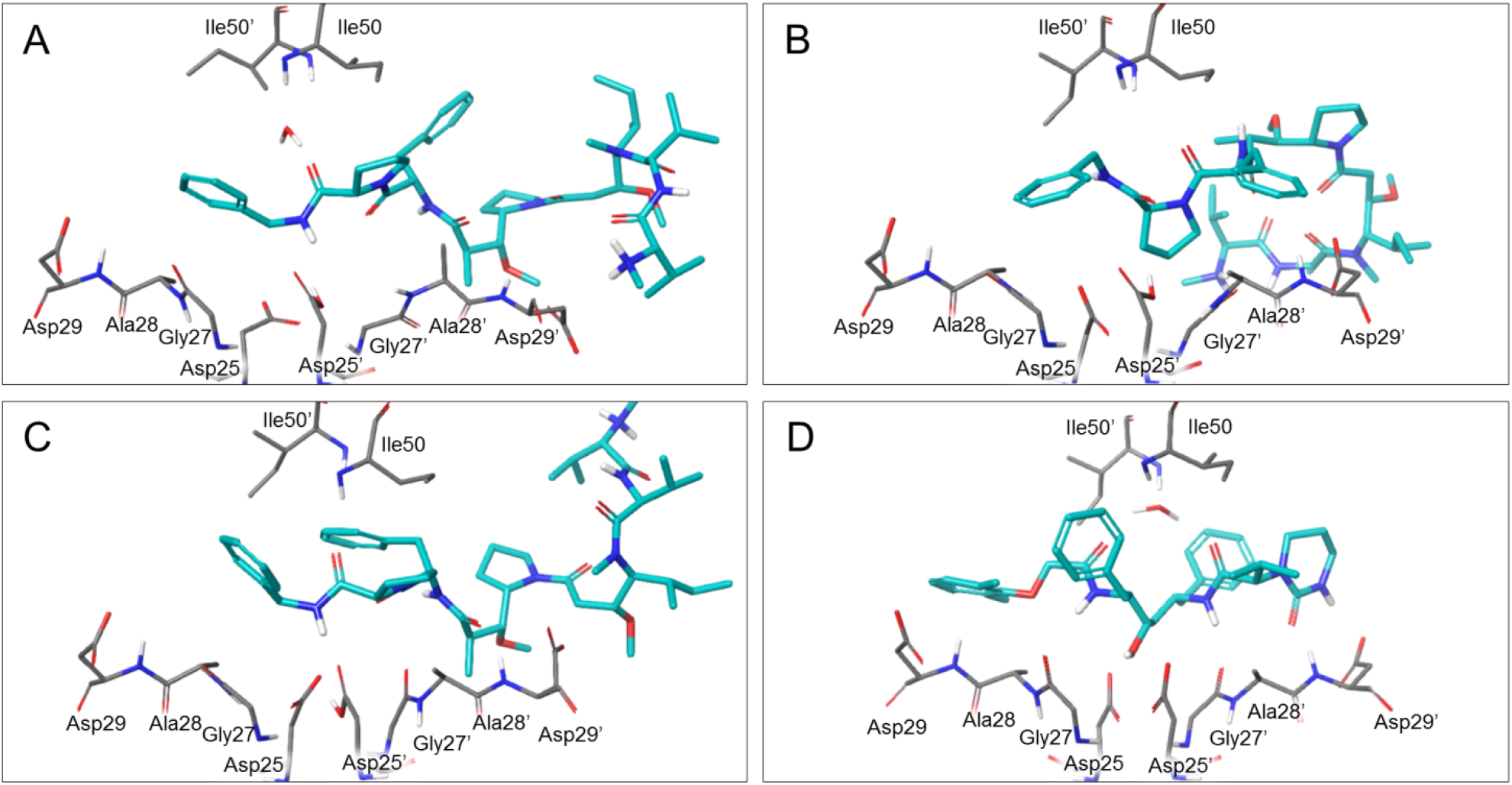
MF-7 binding to HIV-1 protease. A) The most favorable binding mode of MF-7 in HIV-1 protease (SP Gscore = −6.0 kcal/mol) when docked with a bridging water molecule. B) The most favored and C) second most favored binding mode of MF-7 in the protease without inclusion of water molecules in the calculation (SP Gscore = −9.6 and −7.8 kcal/mol, respectively). D) Lopinavir binding to HIV-1 protease as observed in PDB entry 6DJ1. All binding modes are shown from the same view point to ease comparisons and all differences in residue conformations are due to the induced fit docking protocol. In all panels, select residues from subsite S3 to S3’ are shown in gray, while the ligand is shown in cyan. Residue names including an apostrophe denote residues from chain A, while the remaining are from chain B.

## MATERIAL AND METHODS

All chemicals were purchased from commercial vendors (SigmaAldrich, ApiChem, VWR, Merck, Alfa Aesar, Toronto-Research Company, and Tokyo Chemical Industry) and used as delivered unless otherwise stated. Dry solvents (DCM, acetonitrile (MeCN), tetrahydrofuran (THF), and toluene) were obtained from an MBraun SPS-800 solvent purification system, which utilized aluminium oxide for drying. Dry DMF, TEA, DIPEA, and pyridine were purchased from SigmaAldrich/Merck. Ultrapure water was obtained from a miliQ direct 8 system (Milipore).

Thin layer chromatography (TLC) was performed on Merck Kieselgel 60 F254 and visualized by UV and/or stain by submersion into a solution of potassium permanganate, iodine, or ninhydrin followed by blow-drying with heating. Flash column chromatography was performed using silica gel (230-400 mesh particle size, 60 Å pore size) as the stationary phase.

Preparative HPLC was performed on a Gilson HPLC system with a C18 column (Phenomenex, Luna, 5μ, 100Å, 250×10mm) in MeCN/water with UV detection at 254 nm and flow rate of 7mL/min.

NMR spectroscopy was performed using either a Varian Mercury 400 MHz spectrometer or a Bruker Ascend 400 spectrometer, both running at 400 MHz and 101 MHz for ^1^H NMR and ^13^C-NMR respectively. The chemical shifts (δ ppm) were determined using the residual solvent signal as reference. Multiplicities are indicated using the following abbreviations: s = singlet, d = doublet, t = triplet, q = quartet, m = multiplet, bs = broad singlet.

Analytical HPLC was performed on a Schimadzu-LC-2010A system with a C18 column (Ascentris Express Peptide Es-C18, Supelco Analytical) with the following dimension (2.7 m particles, length 150 mm, diameter 3.0 mm). Detection was performed by UV at two wavelengths per run. The mobile phase was a mixture of ultrapure water and MeCN, both of which contained 0.1 % trifluoroacetic acid (TFA) (v/v %).

Mass spectra (High Resolution Mass Spectrometry – HR-MS) were recorded on a Bruker Maxis Impact LC-TOF spectrometer with positive or negative electrospray ionization (ESI). The spectra were recalibrated based on a standard co-injected with the sample.

## ORGANIC SYNTHESES

### Synthesis of Fmoc-MMAF

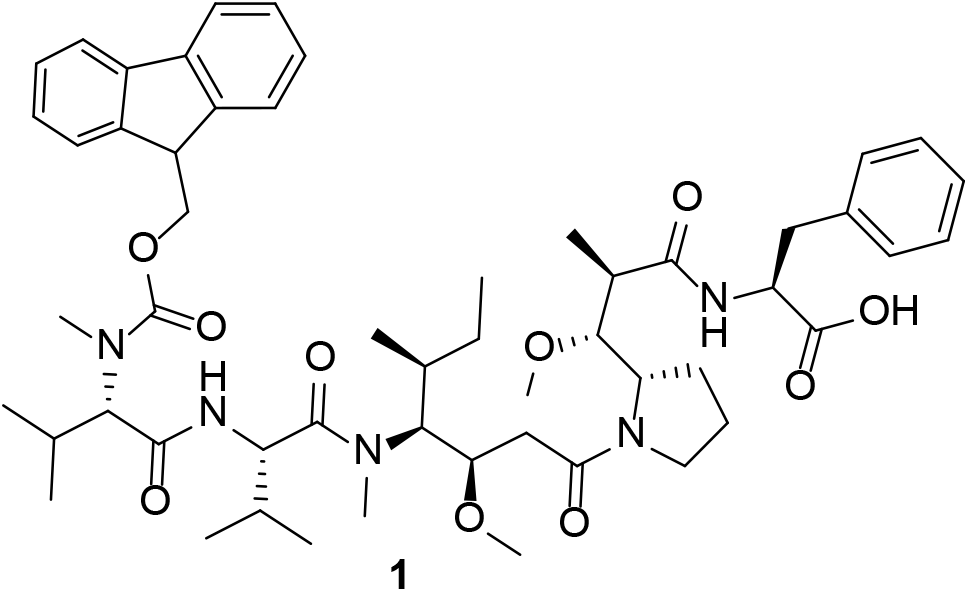

MMAF (40.0 mg, 0.0547 mmol, 1 equiv.) was dissolved in DMF (2 mL). FmocCl (104 mg, 0.401 mmol, 7.3 equiv.) and DIPEA (0.0288 mL, 0.168 mmol, 3 equiv.) were added to the solution sequentially. The reaction was stirred at room temperature for 3 days. The reaction mixture was concentrated and purified by flash column chromatography in MeOH/DCM 0:100 to 2:98 yielding the pure product (52 mg, 0.0547 mmol, 98 %).

**^1^H NMR** (400 MHz, CDCl_3_) δ (ppm) 7.86 −7.70 (m, 3H), 7.63 – 7.47 (m, 3H), 7.44-7.35 (m, 3H), 7.30 (t, *J* = 7.5 Hz, 3H), 7.26-7.19 (m, 3H), 7.15-6.95 (m, *J* = 24.6, 13.1, 4.3 Hz, 1H), 5.09 – 3.77 (m, 7H), 3.50 – 3.19 (m, 5H), 3.17 – 2.68 (m, 6H), 2.53 – 2.19 (m, 2H), 2.09 – 1.59 (m, 4H), 1.19 – 0.41 (m, 18H).

**R**_**f**_ (5:95 MeOH/DCM) = 0.28

### Synthesis of FmocMMAF-Pro-tBu (2)

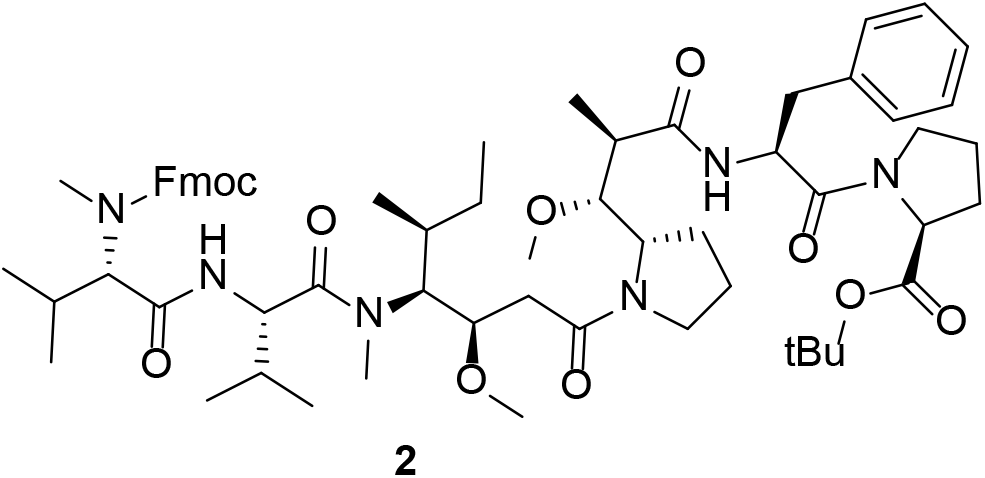

FmocMMAF (**1**) (52 mg, 0.0545 mmol, 1 equiv.) was dissolved in DMF and cooled to 0 °C. DIPEA (0.0559 mL, 0.327 mmol, 6 equiv.) was added followed by *tert*-butylproline (10.3 mg, 0.06 mmol, 1.1 equiv.). HATU (22.8 mg, 0.06 mmol, 1.1 equiv.) was dissolved in DMF (0.5 mL) and added to the reaction mixture over 10 min. The reaction was allowed to heat to room temperature and stirred overnight. The reaction was diluted with saturated sodium bicarbonate and extracted with DCM thrice. The organic phase was dried over MgSO_4_, filtered, and concentrated. The concentrate was purified by flash column chromatography MeOH/DCM (0:100 to 5:95) yielding the pure product (10.0 mg, 0.009 mmol, 17 %).

**^1^H NMR** (400 MHz, CDCl_3_) δ (ppm) 7.86-7.69 (m, 2H), 7.63-7.45 (m, 2H), 7.40 (t, *J* = 7.5 Hz, 2H), 7.30 (t, *J* = 7.5 Hz, 1H), 7.25-6.96 (m, 3H), 5.02 – 4.55 (m, 2H), 4.49-4.34 (m, 2H), 4.26 – 4.09 (m, 2H), 3.86 – 3.24 (m, 8H), 2.80 (s, 3H), 2.56 – 1.61 (m, 16H), 1.57 – 1.41 (m, 9H), 1.26 – 0.52 (m, 19H).

**R**_**f**_ (5:95 MeOH/DCM) = 0.31

### Synthesis of MMAF-Pro-tBu (3)

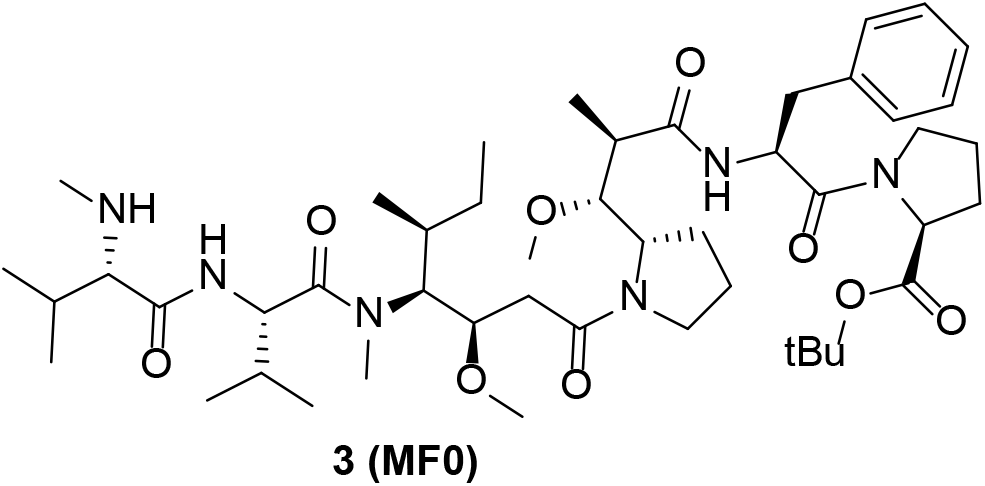

FmocMMAFProtBu (**2**) (10.0 mg, 0.00903 mmol, 1 equiv.) was dissolved in DMF (0.4 mL) and piperidine (0.1 mL), and the reaction stirred at room temperature for 30 min. The reaction mixture was concentrated and purified by trituration with pentane and Et2O yielding the pure product (7.07 mg, 0.00903 mmol, 88 %)

**^1^H NMR** (400 MHz, DMSO-*d6*) δ (ppm) 8.49 – 7.92 (m, 2H), 7.37-7.14 (m, 5H), 4.81-4.36 (m, 3H), 4.19 – 3.50 (m, 6H), 3.23 – 2.64 (m, 26H), 2.38 – 2.18 (m, 7H), 2.08 – 1.50 (m, 19H), 1.50 – 1.29 (m, 9H), 1.13 – 0.65 (m, 24H).

### Synthesis of MMAF-Pro (4)

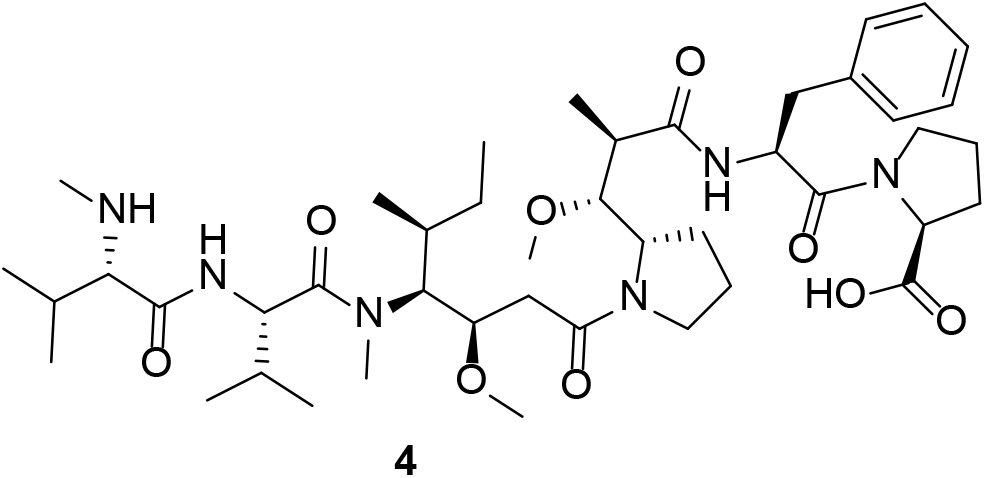

MMAFProtBu (**3**) (3.36 mg, 0.0038 mmol, 1 equiv.) was dissolved in TFA/TIPS/H_2_O cleavage buffer (1 mL, 95:2.5:2.5 v/v) and stirred at room temperature for 2 hours. The reaction was concentrated and purified by trituration in pentane and Et_2_O yielding the pure product. Some solvent was present in pure product as seen in the ^1^H NMR spectrum. The pure product was hence suspended and lyophilized yielding the dry pure product (3.53 mg, 0.0038 mmol, quant.).

**^1^H NMR** (400 MHz, DMSO-*d6*) δ (ppm) 9.03-8.02 (m, 3H), 7.41-7.06 (m, 5H), 4.92-4.36 (m, 4H), 4.25 – 3.45 (m, 6H), 3.27 – 2.73 (m, 17H), 2.43 – 1.33 (m, 17H), 0.94 – 0.58 (m, 21H).

### Synthesis of MMAF-Pro-Gln-Ile (5)

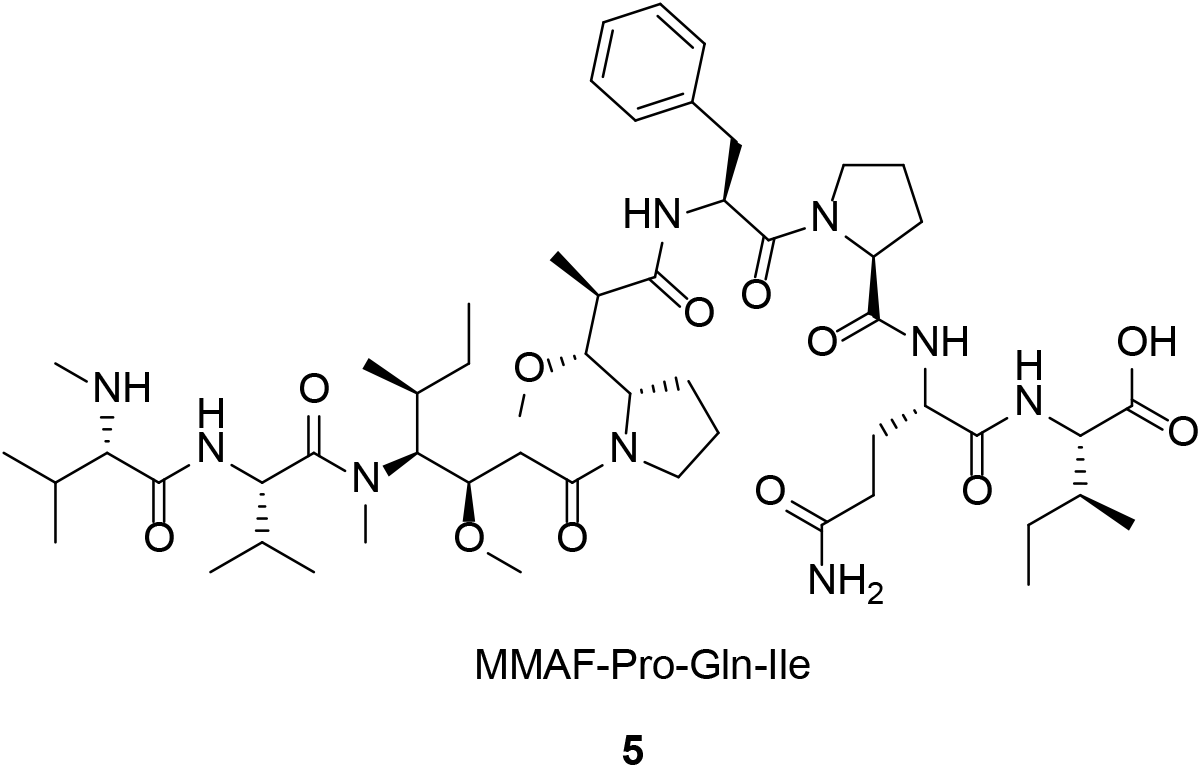

Compound **5** was synthesized through solid phase peptide synthesis using a 2-chlorotrityl (2CT) resin and standard Fmoc protecting group strategy. 2CT resin (1.6 mmol/g, 31 mg, 0.0496 mmol, 1 equiv.) was placed in a syringe. The resin was washed thrice with DMF and thrice with DCM. A solution of FmocIle (22.0 mg, 0.0622 mmol, 1.3 equiv.) and DIPEA (11, μL, 0.0643 mmol, 1.3 equiv.) in DCM (1 mL) was added to the resin and stirred for 5 min. Another aliquot of DIPEA (30 μL, 0.175 mmol, 3.5 equiv.) was added to the reaction, which was stirred for 1 hour at room temperature. Methanol (50 μL) was added to the reaction, which was then stirred for 30 min. The resin was then washed thrice with DMF and thrice with DCM.

The following couplings followed the same pattern. First Fmoc removal by reaction with piperidine/DMF (1:4) for 3 min, 10 min and 10 min. Then a solution of the amino acid (1^st^ FmocGln(Tr)OH (81 mg, 0.133 mmol, 2.7 equiv.), 2^nd^ FmocPro (44.0 mg, 0.13 mmol, 2.6 equiv.)) was activated in DMF with oxyma (13 mg, 0.0915 mmol, 1.8 equiv.) and DIPC (13 μL, 0.0834 mmol, 1.7 equiv.) for 5 min with stirring, upon which it was added to the resin. The resin was stirred at room temperature for 1.5 h and washed thrice with DMF and thrice with DCM. After the coupling, a Kaiser test was performed to confirm coupling to all free amines before deprotection and subsequent coupling.

The last coupling was performed with FmocMMAF (**1**) (52 mg, 0.0545 mmol, 1.1 equiv.) which was pre-activated with oxyma (7 mg, 0.0493 mmol, 1 equiv.) and DIPC (7 μL, 0.0496 mmol, 1 equiv.) in DMF for 5 min. The solution was added to the resin and stirred overnight. Chloroaniline test showed presence of free amines due to incomplete coupling likely due to the low amount of **1** used. The resin was nevertheless washed with DMF thrice and DCM thrice. Fmoc was removed by reaction with piperidine/DMF (1:4) for 3 min, 10 min, and 10 min. The peptide was cleaved from the resin with a 2% TFA in DCM solution (3 × 1 min) followed by washing with DCM thrice. The solvent was removed and the peptide triturated with Et_2_O. ^1^H NMR showed incomplete removal of protecting groups. The peptide was therefore dissolved in 2 mL TFA/H_2_O/DTT (90:5:5) for 1 hour followed by concentration and trituration. Crude MS detected an impurity of Pro-Gln-Ile with no MMAF attached. The crude was purified by preparative HPLC yielding the pure product (27 mg, 0.025 mmol, 51%).

**HR-MS** (ESI) [C_55_H_91_N_9_O_12_+H]^+^ calcd. 1070.6870 found 1070.6857.

#### One-pot preparation of MF1-7 peptides

**Scheme S1.**
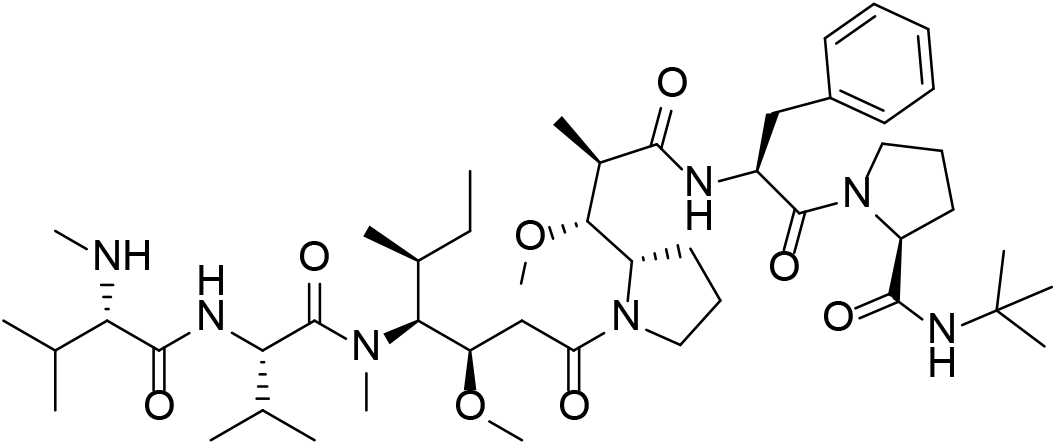
One-pot preparation of **MF1-7**

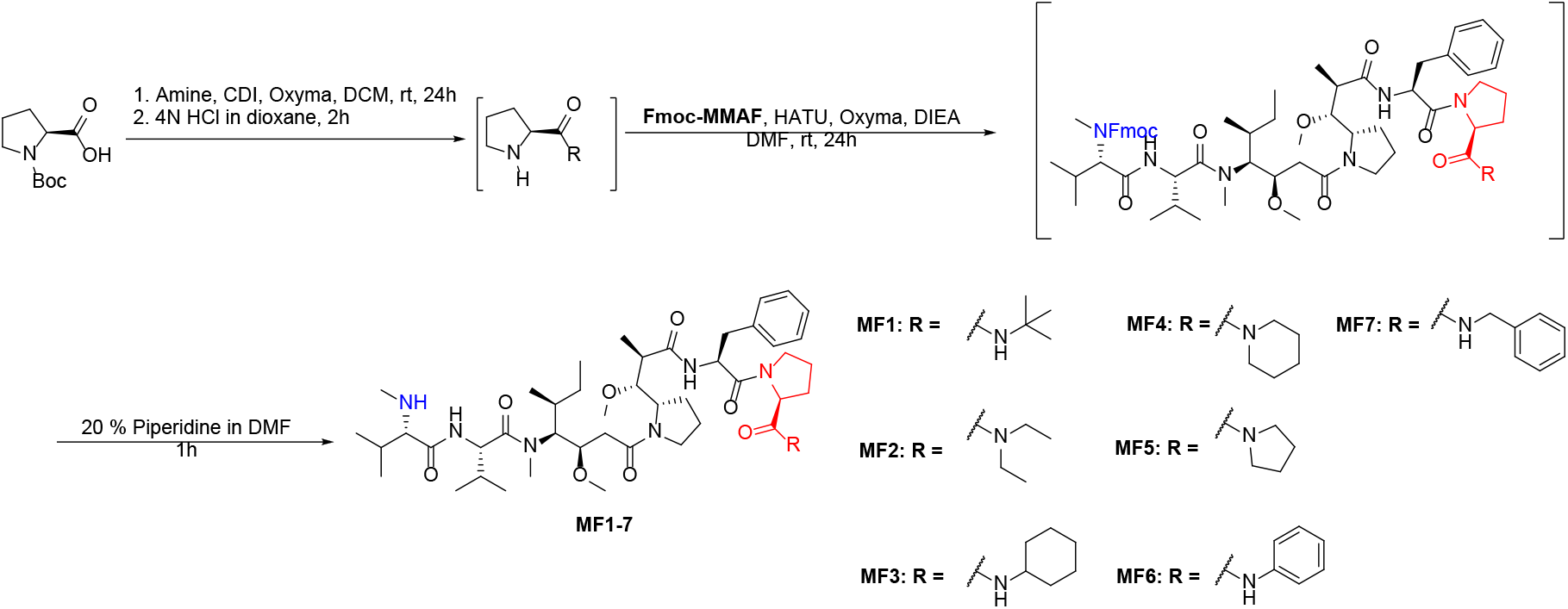

##### MF1

To a solution of Boc-*L*-Pro-OH (10.00 mg, 0.05 mmol, 1 eq.) in DCM, CDI (9.040 mg, 0.056 mmol, 1.2 eq.), Oxyma Pure (13.210 mg, 0.093 mmol, 2 eq.) was added and stirred at room temperature for 5 min. Tert-butyl amine (4.079 mg, 5.86 μL, 0.056 mmol, 1.2 eq.) was then added dropwise. The mixture was stirred at room temperature for 4 h. After fully conversion of Boc-*L*-Pro-OH, the reaction mixture was added to DCM (5 mL) in separatory funnel. The organic layer was washed with 5 mL of NaHCO_3_(sat.) then water (5 mL) and 5 mL of 0.5 % HCl. Organic layer was evaporated to obtain crude product. Dioxane (1 mL) was added to the crude product, followed by 4N HCl in dioxane (1 ml) and resulting was stirred at room temperature for 4h. The reaction mixture was then concentrated and dried for 24h to afford crude free amine.

**Fmoc-MMAF** (6 mg, 0.0062 mmol, 1 eq.), HATU (9.5 mg, 0.025 mmol, 4 eq.), Oxyma Pure (1.8 mg, 0.0124 mmol, 4 eq.) was dissolved in 1 ml of 20 % of collidine. The mixture was then added to a solution of the above-mentioned crude free amine (pre-dissolved in 1 ml 20% collidine in DMF) and stirred for 24h. After consumption of **Fmoc-MMAF** (TLC), the reaction mixture was added to 10 ml of ethyl acetate and washed with 10 mL of 5 % NaHCO_3_ and 10 mL sat. oxalic acid. The organic layer was concentrated to afford crude **Fmoc-MF1**. The crude was washed with cold ethyl ether (3 times), pentane (3 times) and used directly for the final deprotection.

**Figure S3.**
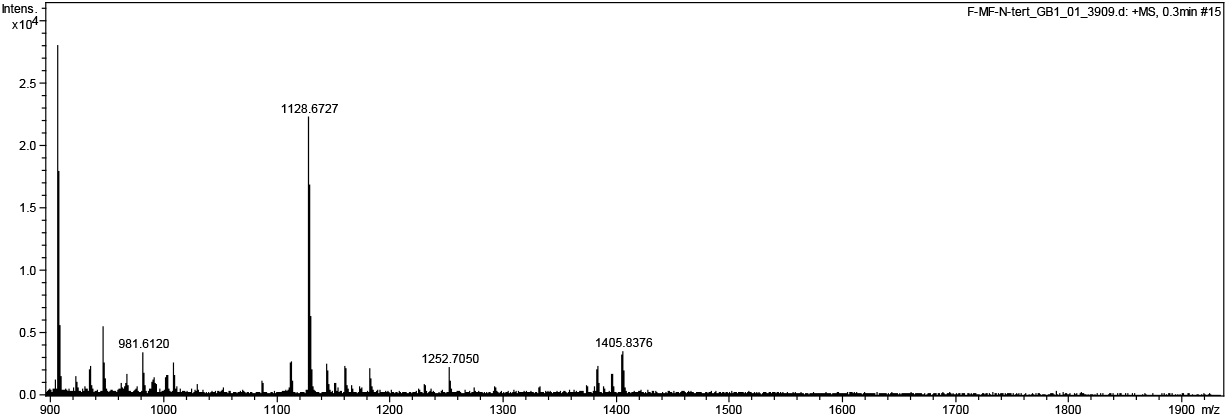
HRMS of curde **Fmoc-MF1** before deprotection. Cal. For C_63_H_91_N_7_NaO_10_ [M+Na]^+^ 1128.6725, found 1128.6727.

To a solution of crude **Fmoc-MF1** in 0.5 ml DMF, 1ml of 20% piperidine in DMF was added and stirred for 5 min. The reaction mixture was concentrated under reduced pressure to afford crude **MF1**. The crude was washed with cold ethyl ether (3 times), pentane (3 times) and purified by preparative-HPLC to afford 3.2 mg of final product.

**Figure S4.**
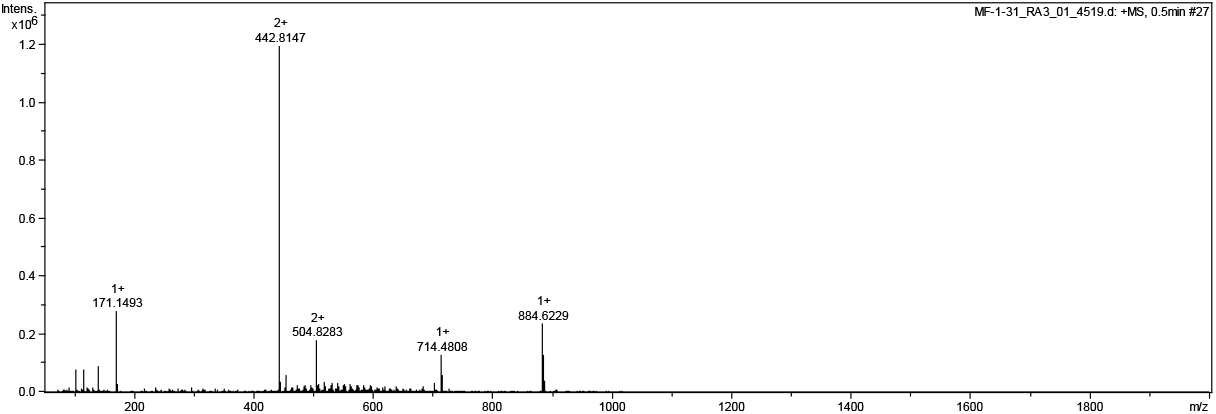
HRMS of **MF1** HRMS (ESI+) Cal. For C_48_H_82_N_7_O_8_ [M+H]^+^ 884.6225 found 884.6229.

**Figure S5.**
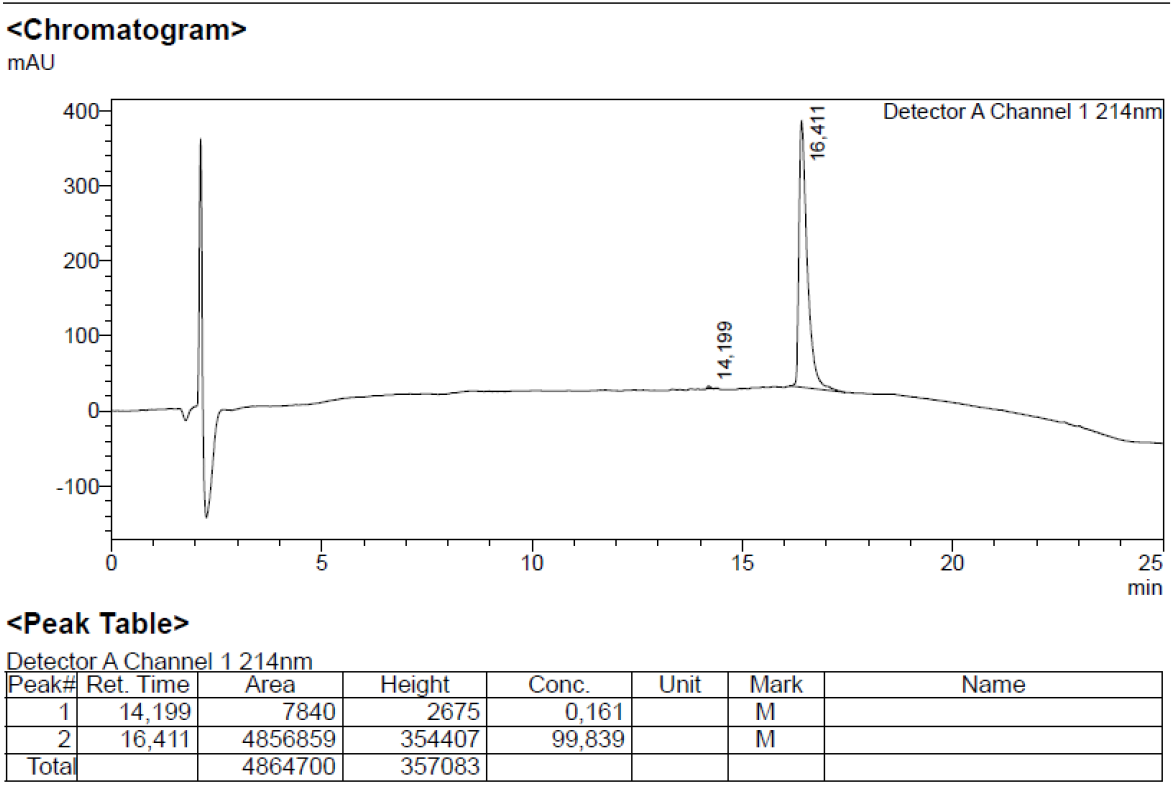
Analytical HPLC of **MF1**.

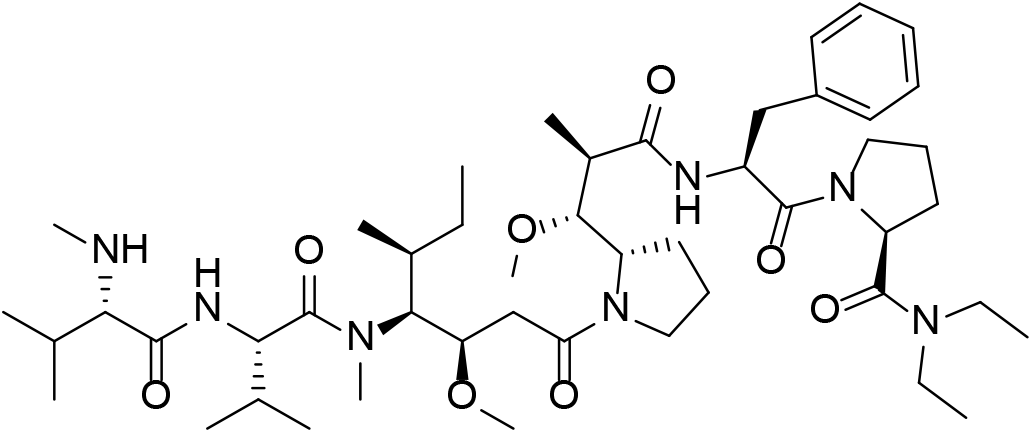

##### MF2

To a solution of Boc-*L*-Pro-OH (10.00 mg, 0.05 mmol, 1 eq.) in DCM, CDI (9.040 mg, 0.056 mmol, 1.2 eq.), Oxyma Pure (13.210 mg, 0.093 mmol, 2 eq.) was added and stirred at room temperature for 5 min. Diethylamine (4.079 mg, 5.77 μL, 0.056 mmol, 1.2 eq.) was then added dropwise. The mixture was stirred at room temperature for 4 h. After fully conversion of Boc-*L*-Pro-OH, the reaction mixture was added to DCM (5 mL) in separatory funnel. The organic layer was washed with 5 mL of NaHCO_3_(sat.) then water (5 mL) and 5 mL of 0.5 % HCl. Organic layer was evaporated to obtain crude product. Dioxane (1 mL) was added to the crude product, followed by 4N HCl in dioxane (1 ml) and resulting was stirred at room temperature for 4h. The reaction mixture was then concentrated and dried for 24h to afford crude free amine.

**Fmoc-MMAF** (6 mg, 0.0062 mmol, 1 eq.), HATU (9.5 mg, 0.025 mmol, 4 eq.), Oxyma Pure (1.8 mg, 0.0124 mmol, 4 eq.) was dissolved in 1 ml of 20 % of collidine. The mixture was then added to a solution of the above-mentioned crude free amine (pre-dissolved in 1 ml 20% collidine in DMF) and stirred for 24h. After consumption of **Fmoc-MMAF** (TLC), the reaction mixture was added to 10 ml of ethyl acetate and washed with 10 mL of 5 % NaHCO_3_ and 10 mL sat. oxalic acid. The organic layer was concentrated to afford crude **Fmoc-MF2**. The crude was washed with cold ethyl ether (3 times), pentane (3 times) and used directly for the final deprotection.

To a solution of crude **Fmoc-MF2** in 0.5 ml DMF, 1ml of 20% piperidine in DMF was added and stirred for 5 min. The reaction mixture was concentrated under reduced pressure to afford crude **MF2**. The crude was washed with cold ethyl ether (3 times), pentane (3 times) and purified by preparative-HPLC to afford 2.8 mg of final product.

HRMS of **MF2** HRMS (ESI+) Cal. For [M+H]^+^ 884.6225 found 884.6229.

**Figure S6.**
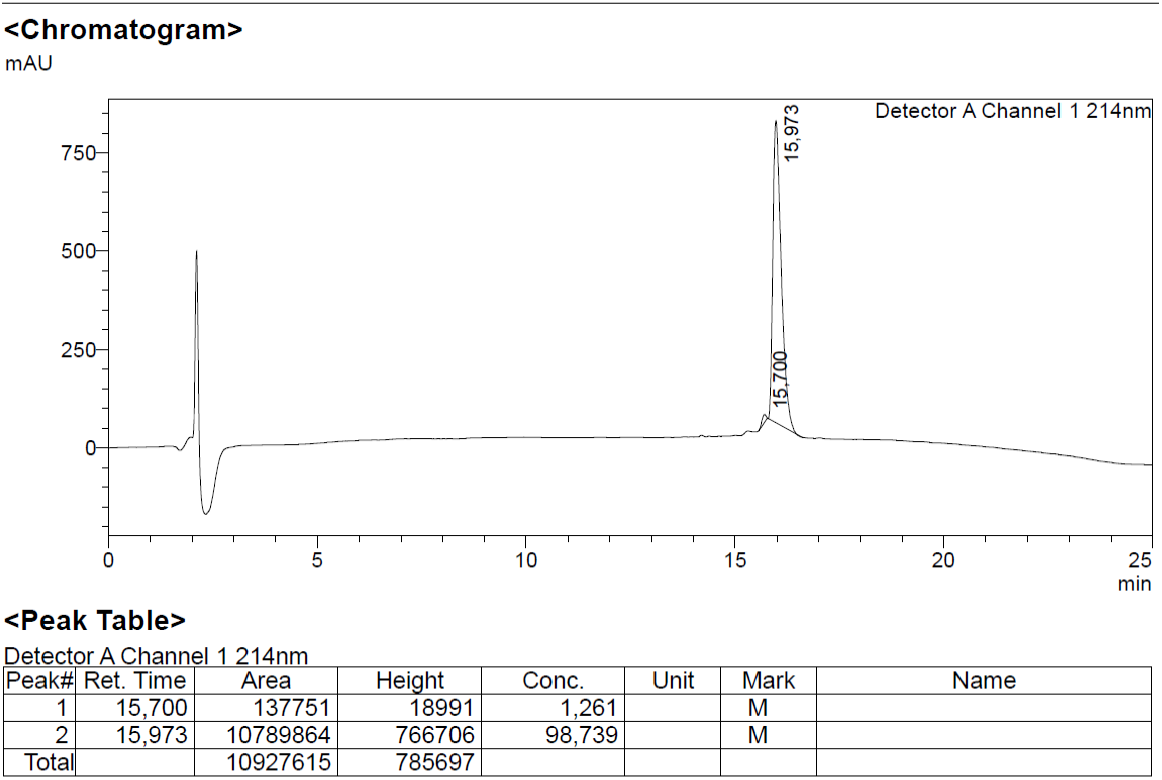
Analytical HPLC of **MF2**.

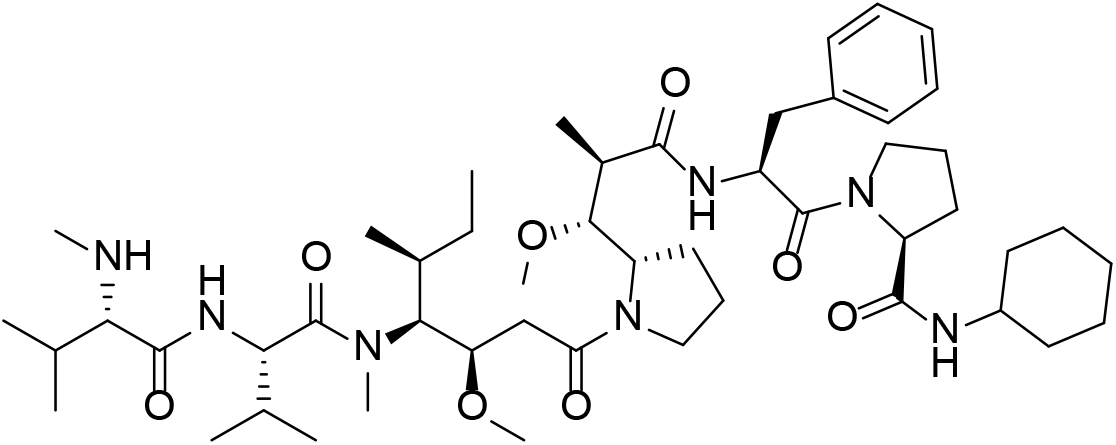

##### MF3

To a solution of Boc-*L*-Pro-OH (10.00 mg, 0.05 mmol, 1 eq.) in DCM, CDI (9.040 mg, 0.056 mmol, 1.2 eq.), Oxyma Pure (13.210 mg, 0.093 mmol, 2 eq.) was added and stirred at room temperature for 5 min. Cyclohexylamine (5.531 mg, 6.40 μL, 0.056 mmol, 1.2 eq.) was then added dropwise. The mixture was stirred at room temperature for 4 h. After fully conversion of Boc-*L*-Pro-OH, the reaction mixture was added to DCM (5 mL) in separatory funnel. The organic layer was washed with 5 mL of NaHCO_3_(sat.) then water (5 mL) and 5 mL of 0.5 % HCl. Organic layer was evaporated to obtain crude product. Dioxane (1 mL) was added to the crude product, followed by 4N HCl in dioxane (1 ml) and resulting was stirred at room temperature for 4h. The reaction mixture was then concentrated and dried for 24h to afford crude free amine.

**Fmoc-MMAF** (6 mg, 0.0062 mmol, 1 eq.), HATU (9.5 mg, 0.025 mmol, 4 eq.), Oxyma Pure (1.8 mg, 0.0124 mmol, 4 eq.) was dissolved in 1 ml of 20 % of collidine. The mixture was then added to a solution of the above-mentioned crude free amine (pre-dissolved in 1 ml 20% collidine in DMF) and stirred for 24h. After consumption of **Fmoc-MMAF** (TLC), the reaction mixture was added to 10 ml of ethyl acetate and washed with 10 mL of 5 % NaHCO_3_ and 10 mL sat. oxalic acid. The organic layer was concentrated to afford crude **Fmoc-MF3**. The crude was washed with cold ethyl ether (3 times), pentane (3 times) and used directly for the final deprotection.

**Figure S7.**
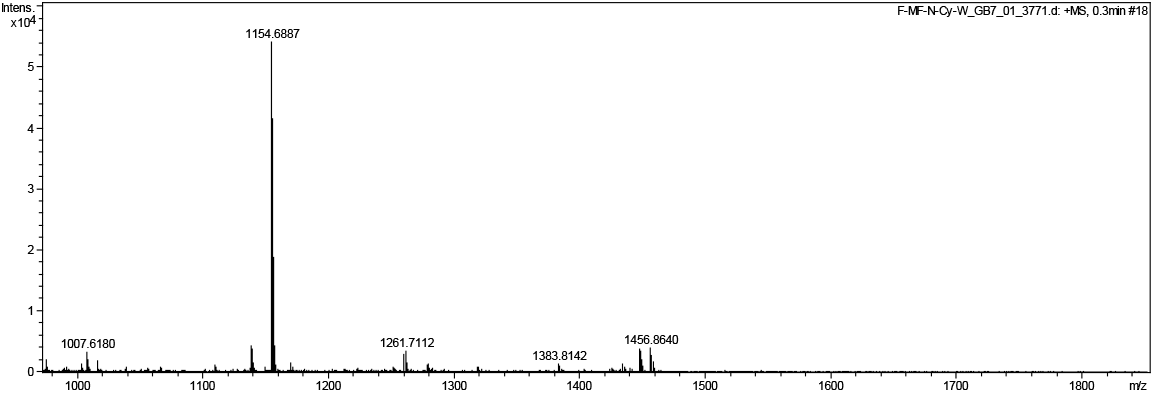
HRMS of crude **Fmoc-MF3** Cal. For [M+Na] ^+^ 1154.6882, found 1154.6887

To a solution of crude **Fmoc-MF3** in 0.5 ml DMF, 1ml of 20% piperidine in DMF was added and stirred for 5 min. The reaction mixture was concentrated under reduced pressure to afford crude **MF3**. The crude was washed with cold ethyl ether (3 times), pentane (3 times) and purified by preparative-HPLC to afford 4.9 mg of final product.

**Figure S8.**
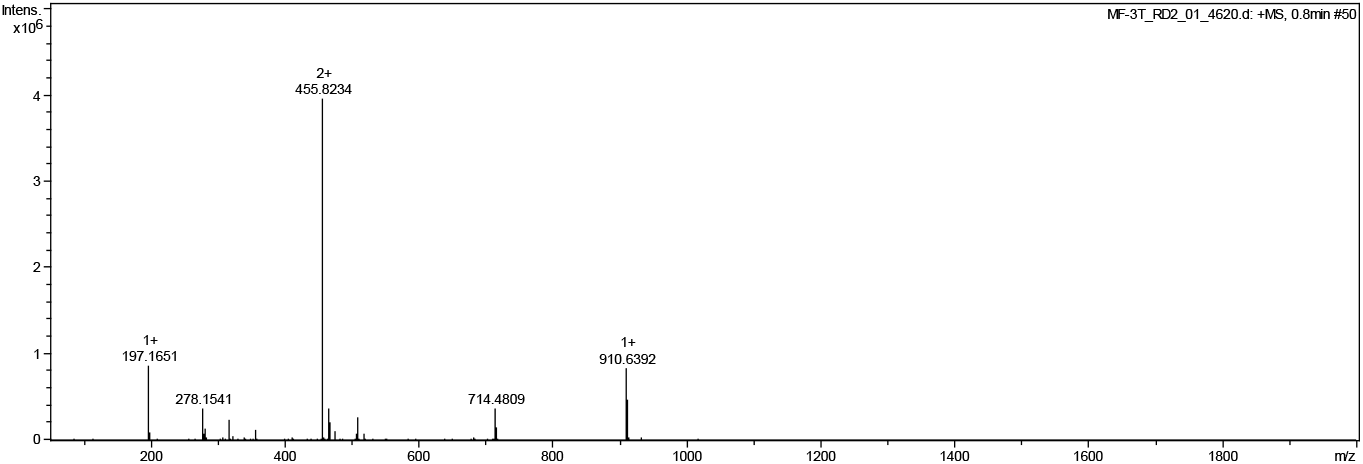
HRMS of **MF3** Cal. For C_50_H_84_N_7_O_8_ [M+H]^+^ 910.6381, found 910.6392.

**Figure S9.**
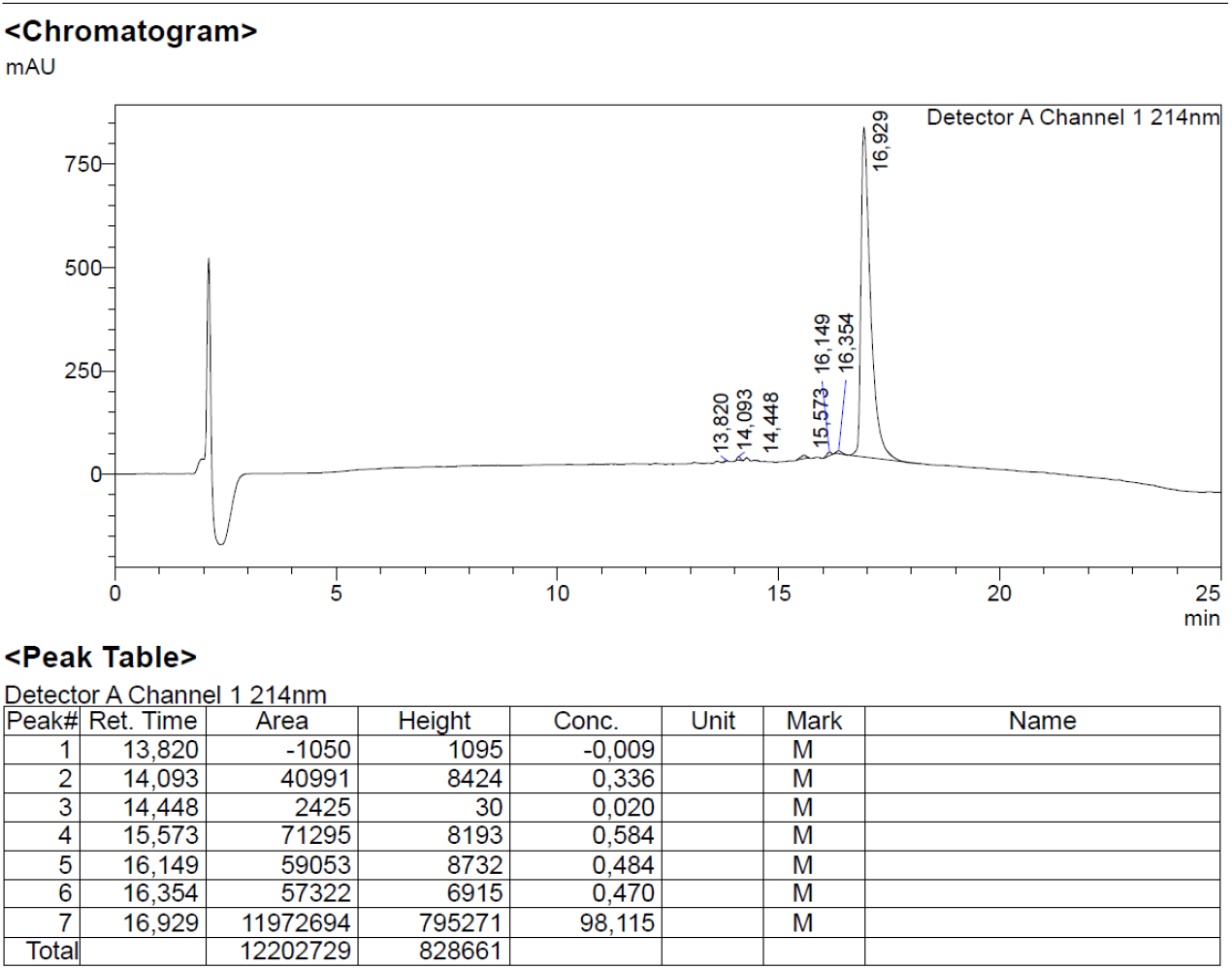
Analytical HPLC of **MF3**.

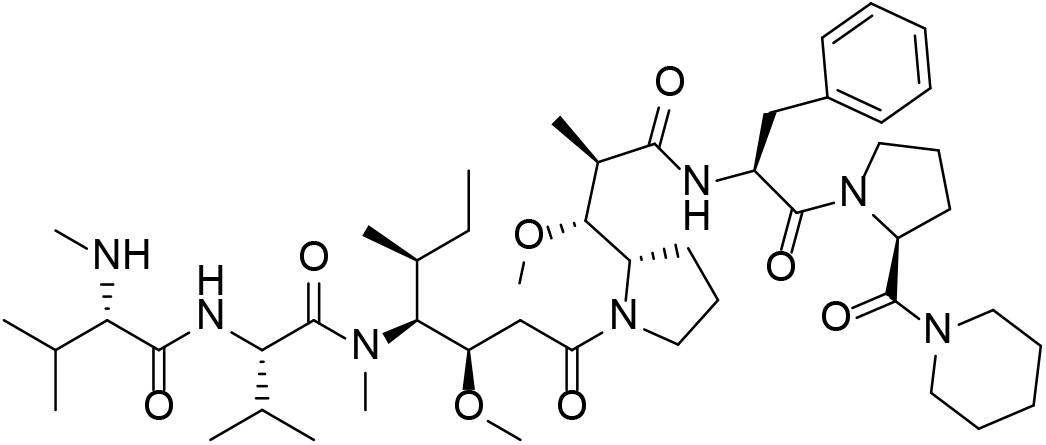

##### MF4

To a solution of Boc-*L*-Pro-OH (10.00 mg, 0.05 mmol, 1 eq.) in DCM, CDI (9.040 mg, 0.056 mmol, 1.2 eq.), Oxyma Pure (13.210 mg, 0.093 mmol, 2 eq.) was added and stirred at room temperature for 5 min. Piperidine (4.749 mg, 5.51 μL, 0.056 mmol, 1.2 eq.) was then added dropwise. The mixture was stirred at room temperature for 4 h. After fully conversion of Boc-*L*-Pro-OH, the reaction mixture was added to DCM (5 mL) in separatory funnel. The organic layer was washed with 5 mL of NaHCO_3_(sat.) then water (5 mL) and 5 mL of 0.5 % HCl. Organic layer was evaporated to obtain crude product. Dioxane (1 mL) was added to the crude product, followed by 4N HCl in dioxane (1 ml) and resulting was stirred at room temperature for 4h. The reaction mixture was then concentrated and dried for 24h to afford crude free amine.

**Fmoc-MMAF** (6 mg, 0.0062 mmol, 1 eq.), HATU (9.5 mg, 0.025 mmol, 4 eq.), Oxyma Pure (1.8 mg, 0.0124 mmol, 4 eq.) was dissolved in 1 ml of 20 % of collidine. The mixture was then added to a solution of the above-mentioned crude free amine (pre-dissolved in 1 ml 20% collidine in DMF) and stirred for 24h. After consumption of **Fmoc-MMAF** (TLC), the reaction mixture was added to 10 ml of ethyl acetate and washed with 10 mL of 5 % NaHCO_3_ and 10 mL sat. oxalic acid. The organic layer was concentrated to afford crude **Fmoc-MF4**. The crude was washed with cold ethyl ether (3 times), pentane (3 times) and used directly for the final deprotection.

**Figure S10.**
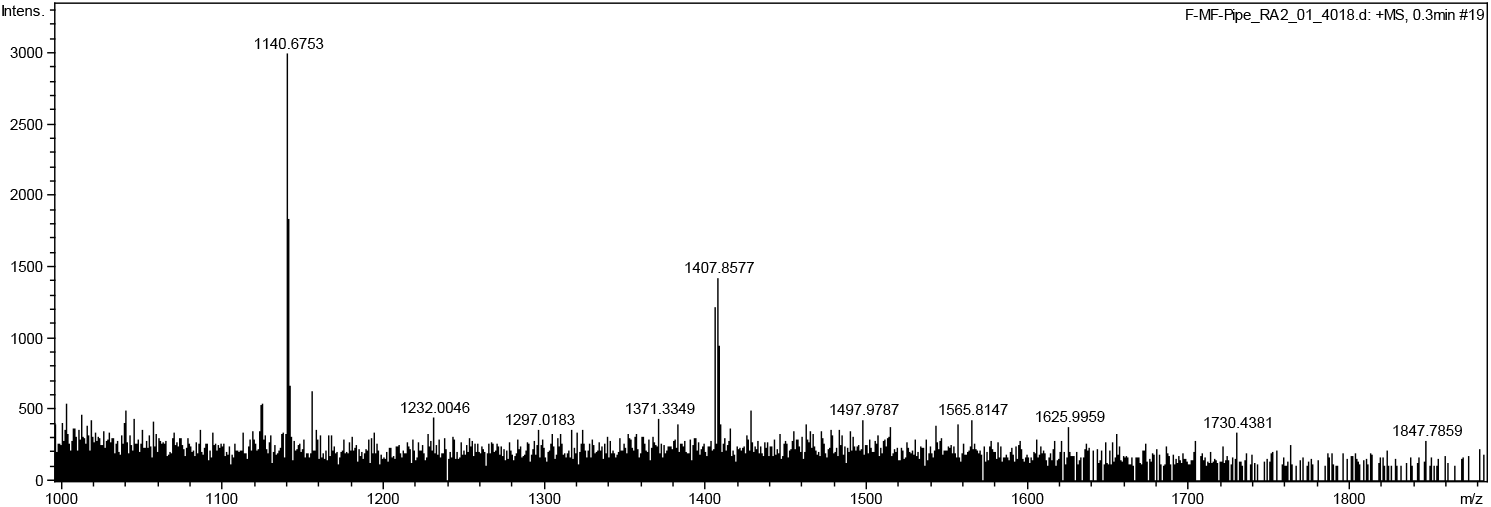
HRMS of crude **Fmoc-MF4** Cal. For [M+Na]^+^ 1140.6725, found 1140.6753

To a solution of crude **Fmoc-MF4** in 0.5 ml DMF, 1ml of 20% piperidine in DMF was added and stirred for 5 min. The reaction mixture was concentrated under reduced pressure to afford crude **MF4**. The crude was washed with cold ethyl ether (3 times), pentane (3 times) and purified by preparative-HPLC to afford 2.7 mg of final product.

**Figure S11.**
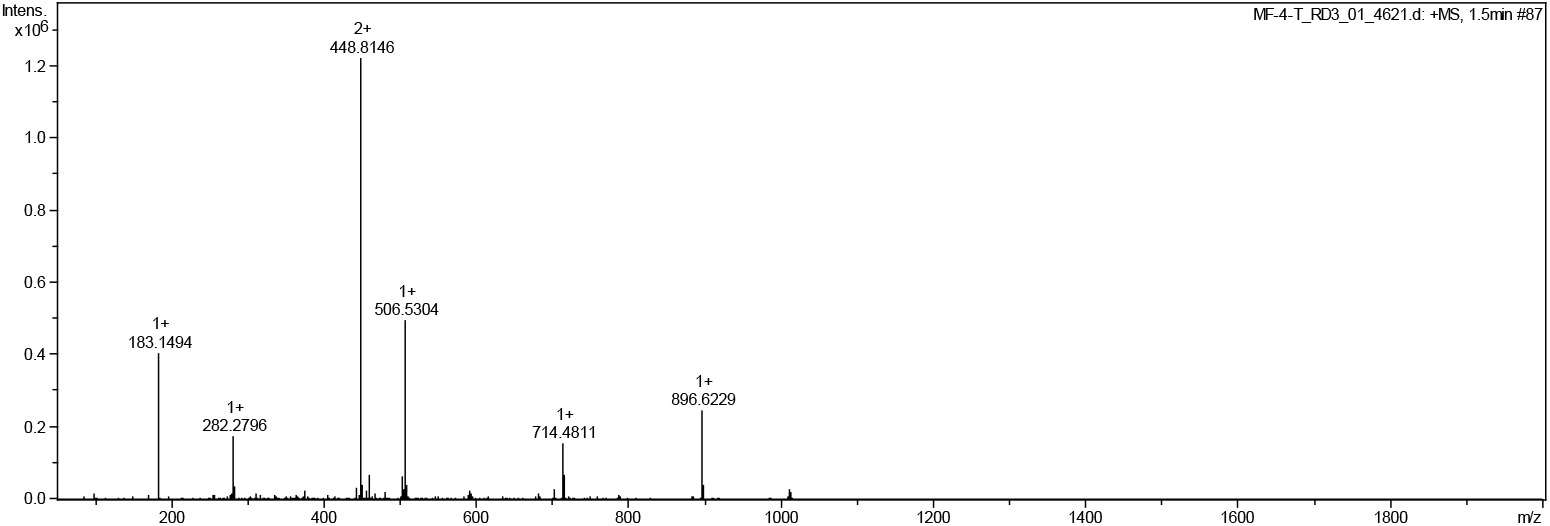
HRMS of **MF4** Cal. For C_49_H_82_N_7_O_8_ [M+H]^+^ 896.6225, found 896.6229.

**Figure S10.**
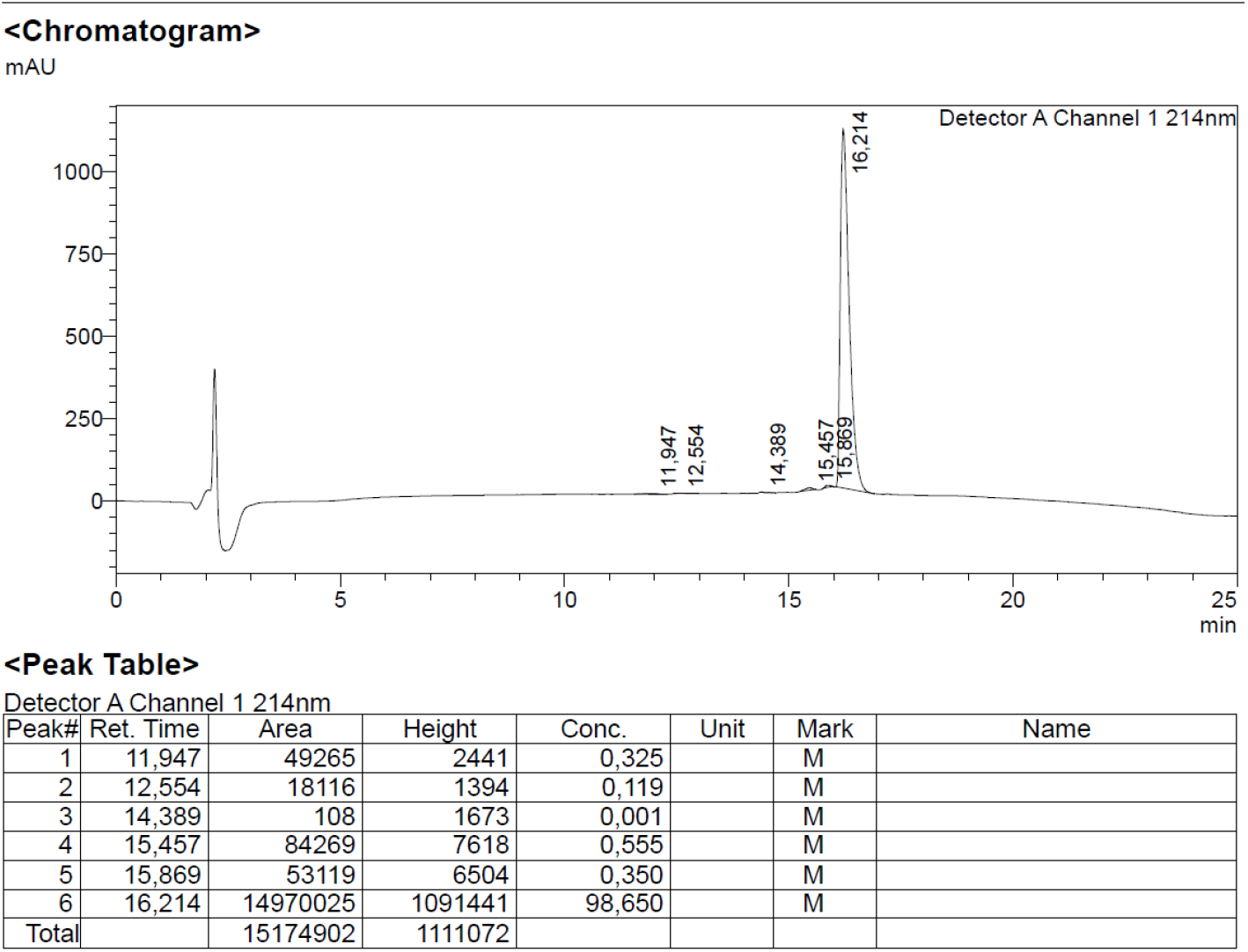
Analytical HPLC of **MF4**

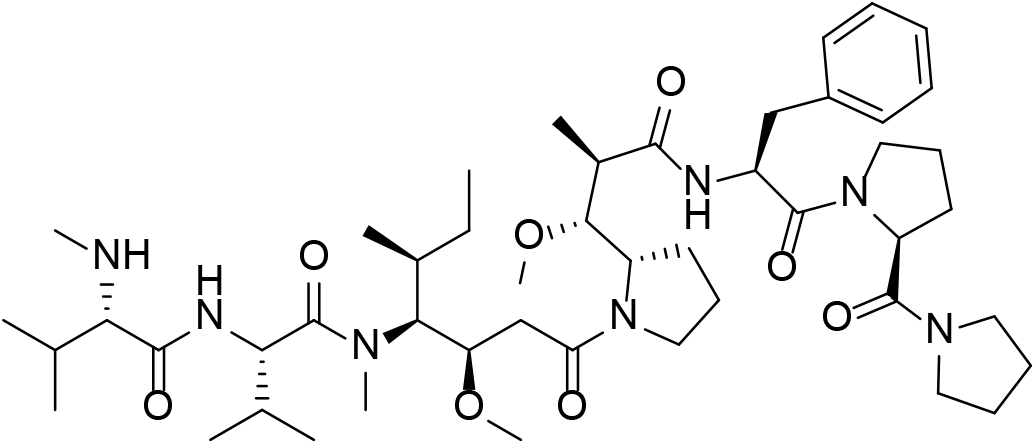

##### MF5

To a solution of Boc-*L*-Pro-OH (10.00 mg, 0.05 mmol, 1 eq.) in DCM, CDI (9.040 mg, 0.056 mmol, 1.2 eq.), Oxyma Pure (13.210 mg, 0.093 mmol, 2 eq.) was added and stirred at room temperature for 5 min. Pyrrolidine (3.967 mg, 4.58 μL, 0.056 mmol, 1.2 eq.) was then added dropwise. The mixture was stirred at room temperature for 4 h. After fully conversion of Boc-*L*-Pro-OH, the reaction mixture was added to DCM (5 mL) in separatory funnel. The organic layer was washed with 5 mL of NaHCO_3_(sat.) then water (5 mL) and 5 mL of 0.5 % HCl. Organic layer was evaporated to obtain crude product. Dioxane (1 mL) was added to the crude product, followed by 4N HCl in dioxane (1 ml) and resulting was stirred at room temperature for 4h. The reaction mixture was then concentrated and dried for 24h to afford crude free amine.

**Fmoc-MMAF** (6 mg, 0.0062 mmol, 1 eq.), HATU (9.5 mg, 0.025 mmol, 4 eq.), Oxyma Pure (1.8 mg, 0.0124 mmol, 4 eq.) was dissolved in 1 ml of 20 % of collidine. The mixture was then added to a solution of the above-mentioned crude free amine (pre-dissolved in 1 ml 20% collidine in DMF) and stirred for 24h. After consumption of **Fmoc-MMAF** (TLC), the reaction mixture was added to 10 ml of ethyl acetate and washed with 10 mL of 5 % NaHCO_3_ and 10 mL sat. oxalic acid. The organic layer was concentrated to afford crude **Fmoc-MF5**. The crude was washed with cold ethyl ether (3 times), pentane (3 times) and used directly for the final deprotection.

**Figure S13.**
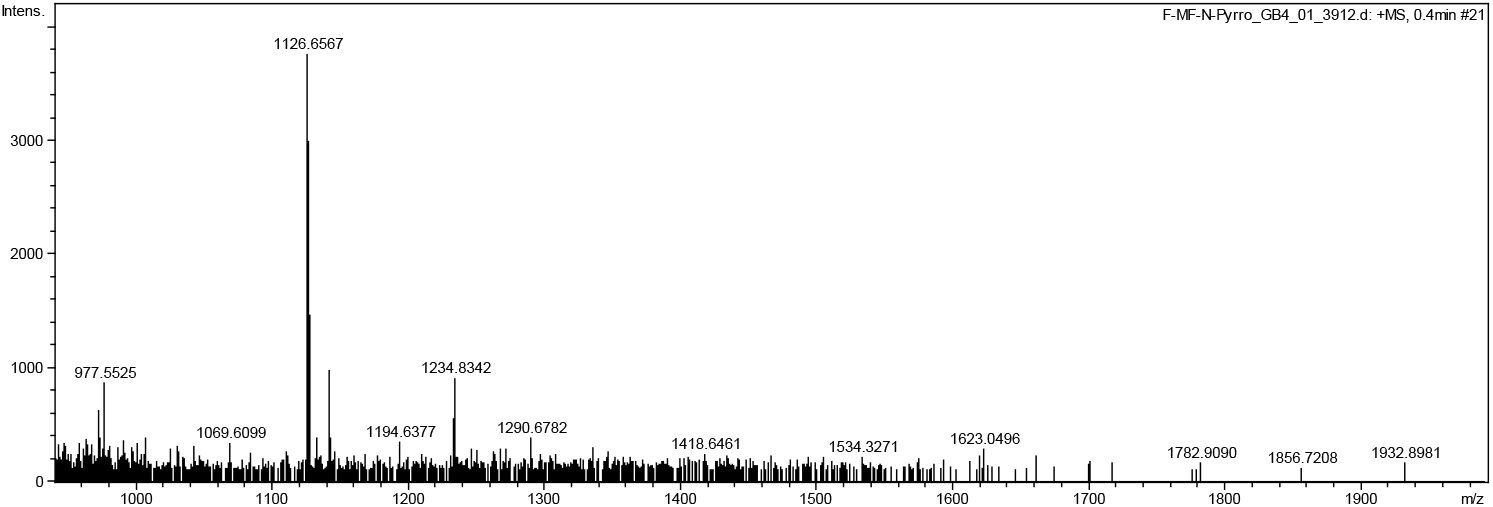
HRMS of crude **Fmoc-MF5** Cal. For [M+Na] ^+^ 1126.6568, found 1126.6567

To a solution of crude **Fmoc-MF5** in 0.5 ml DMF, 1ml of 20% piperidine in DMF was added and stirred for 5 min. The reaction mixture was concentrated under reduced pressure to afford crude **MF5**. The crude was washed with cold ethyl ether (3 times), pentane (3 times) and purified by preparative-HPLC to afford 5.7 mg of final product.

**Figure S14.**
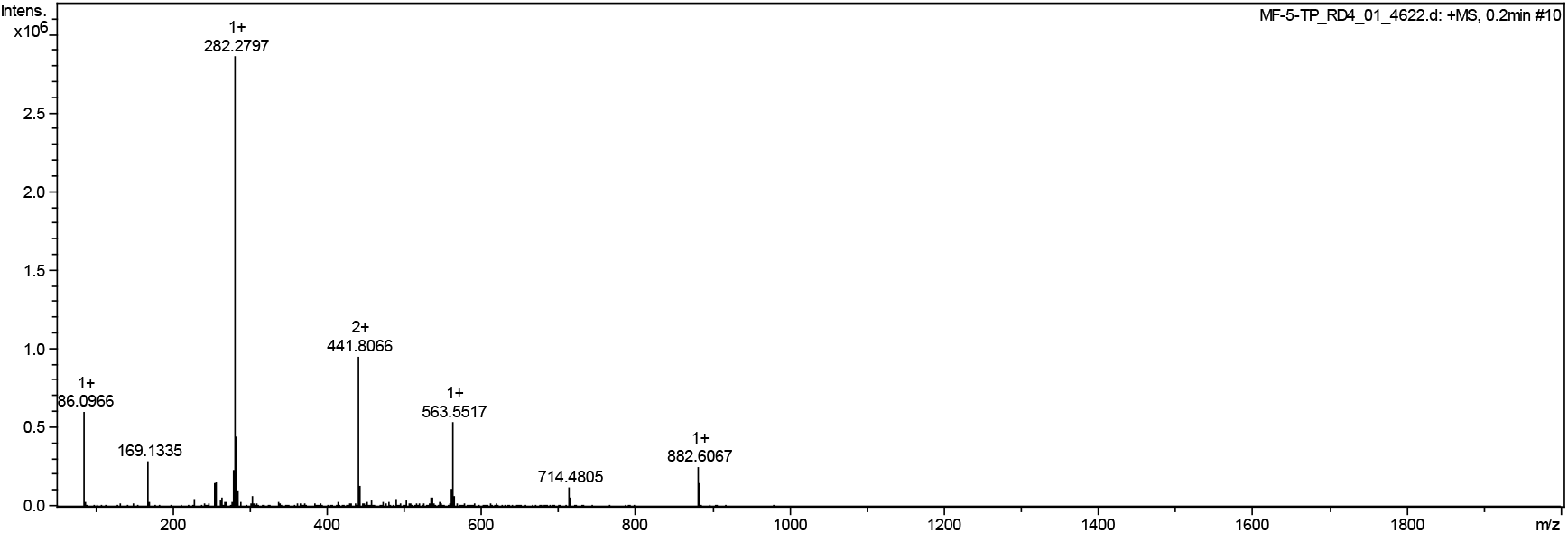
HRMS of **MF5** Cal. For: C_48_H_80_N_7_O_8_ [M+H]^+^ 882.6068, found 882.6067

**Figure S13.**
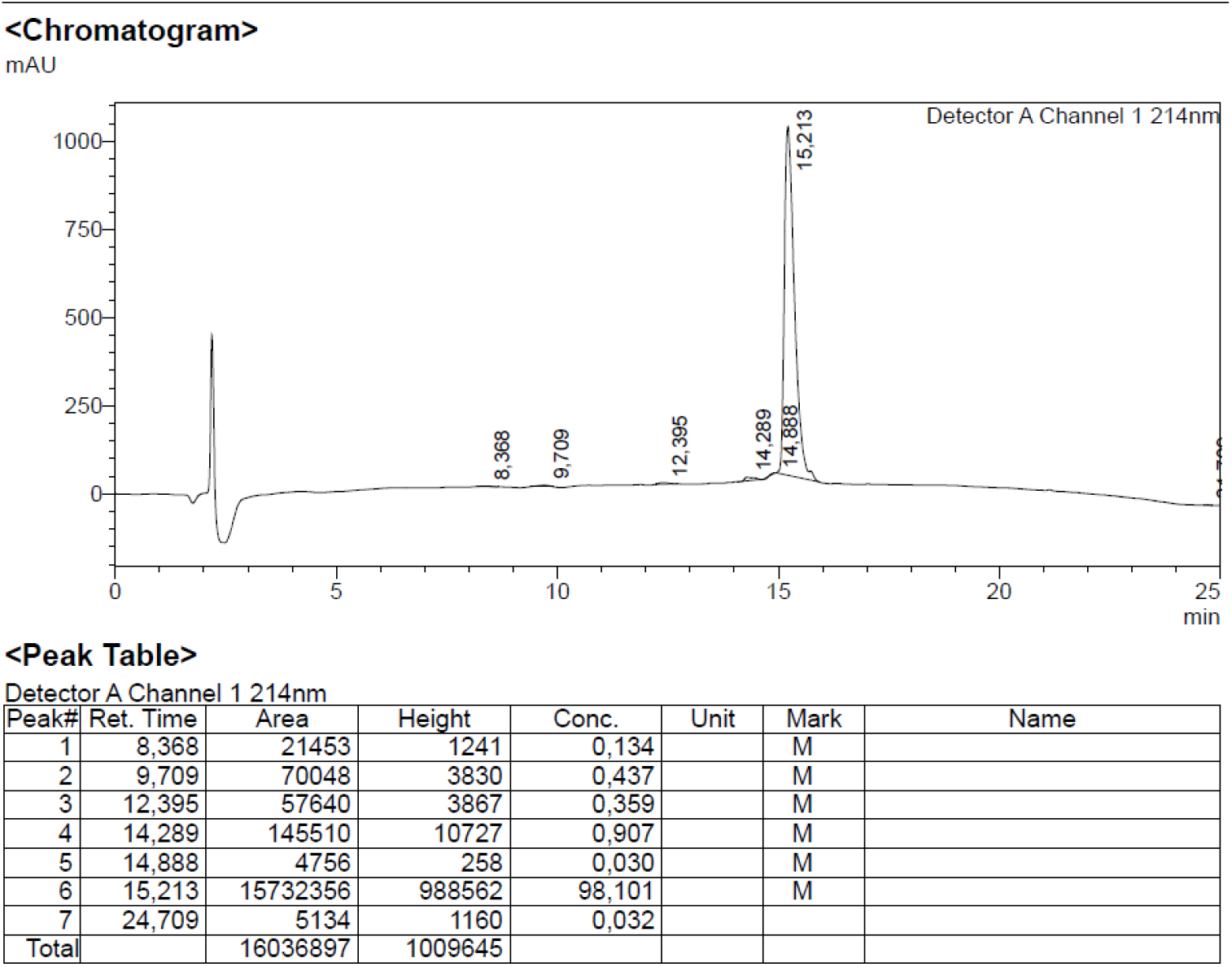
Analytical HPLC of **MF5**

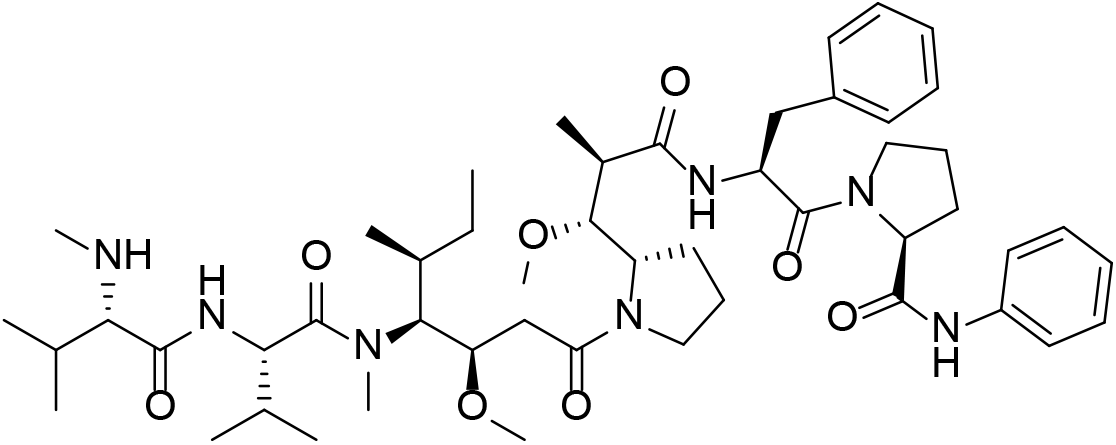

##### MF6

To a solution of Boc-*L*-Pro-OH (10.00 mg, 0.05 mmol, 1 eq.) in DCM, CDI (9.040 mg, 0.056 mmol, 1.2 eq.), Oxyma Pure (13.210 mg, 0.093 mmol, 2 eq.) was added and stirred at room temperature for 5 min. Aniline (5.194 mg, 5.04 μL, 0.056 mmol, 1.2 eq.) was then added dropwise. The mixture was stirred at room temperature for 4 h. After fully conversion of Boc-*L*-Pro-OH, the reaction mixture was added to DCM (5 mL) in separatory funnel. The organic layer was washed with 5 mL of NaHCO_3_(sat.) then water (5 mL) and 5 mL of 0.5 % HCl. Organic layer was evaporated to obtain crude product. Dioxane (1 mL) was added to the crude product, followed by 4N HCl in dioxane (1 ml) and resulting was stirred at room temperature for 4h. The reaction mixture was then concentrated and dried for 24h to afford crude free amine.

**Fmoc-MMAF** (6 mg, 0.0062 mmol, 1 eq.), HATU (9.5 mg, 0.025 mmol, 4 eq.), Oxyma Pure (1.8 mg, 0.0124 mmol, 4 eq.) was dissolved in 1 ml of 20 % of collidine. The mixture was then added to a solution of the above-mentioned crude free amine (pre-dissolved in 1 ml 20% collidine in DMF) and stirred for 24h. After consumption of **Fmoc-MMAF** (TLC), the reaction mixture was added to 10 ml of ethyl acetate and washed with 10 mL of 5 % NaHCO_3_ and 10 mL sat. oxalic acid. The organic layer was concentrated to afford crude **Fmoc-MF6**. The crude was washed with cold ethyl ether (3 times), pentane (3 times) pentane and used directly for the final deprotection.

**Figure S16.**
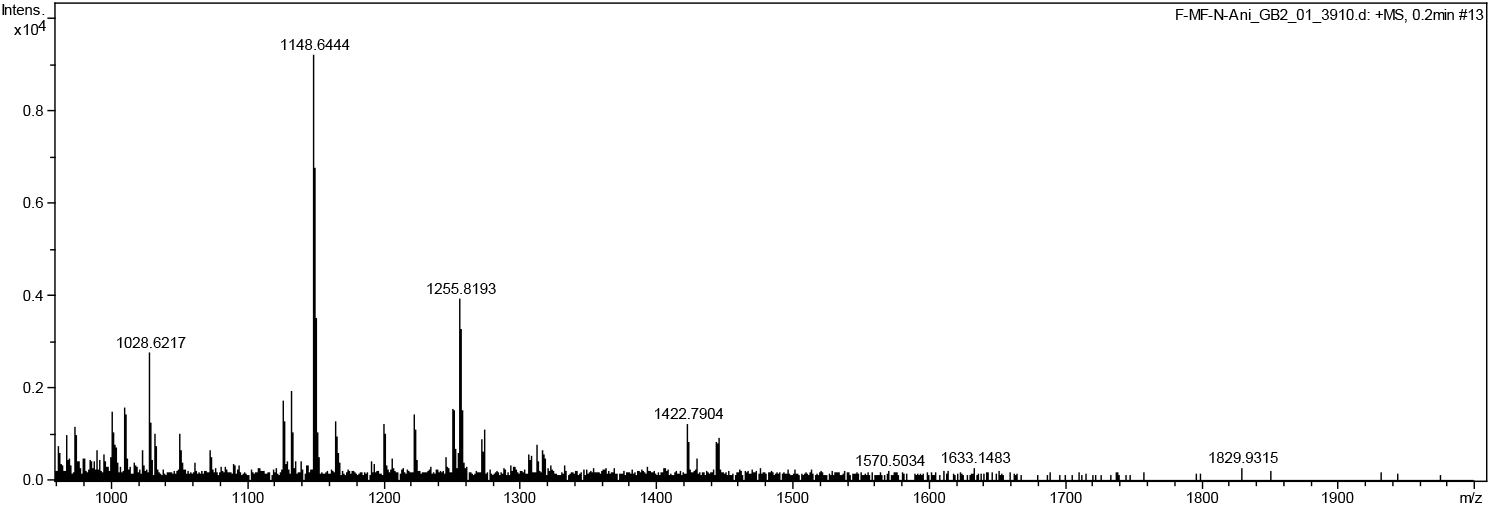
HRMS of crude **Fmoc-MF6** Cal. For [M+Na]^+^ 1148.6412, found 1148.6444

To a solution of crude **Fmoc-MF6** in 0.5 ml DMF, 1ml of 20% piperidine in DMF was added and stirred for 5 min. The reaction mixture was concentrated under reduced pressure to afford crude **MF6**. The crude was washed with cold ethyl ether (3 times), pentane (3 times) and purified by preparative-HPLC to afford 5.1 mg of final product.

**Figure S17.**
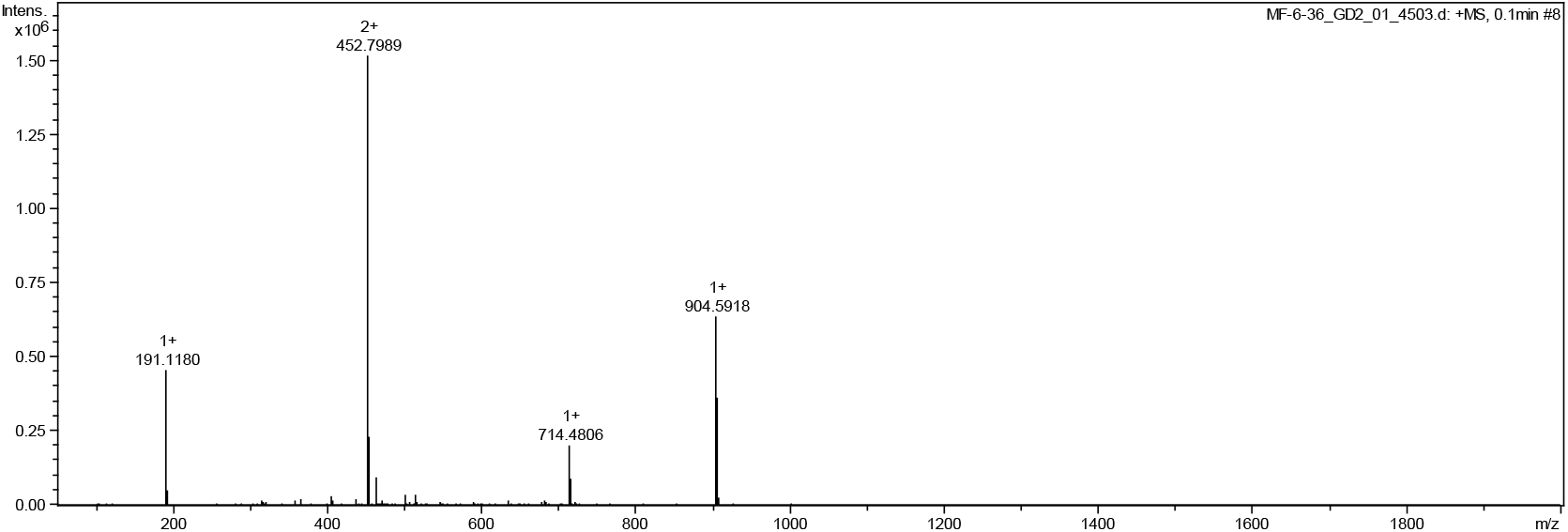
HRMS of **MF6** Cal. For C_50_H_78_N_7_O_8_ [M+H]^+^ 904.5912, found 904.5918

**Figure S16.**
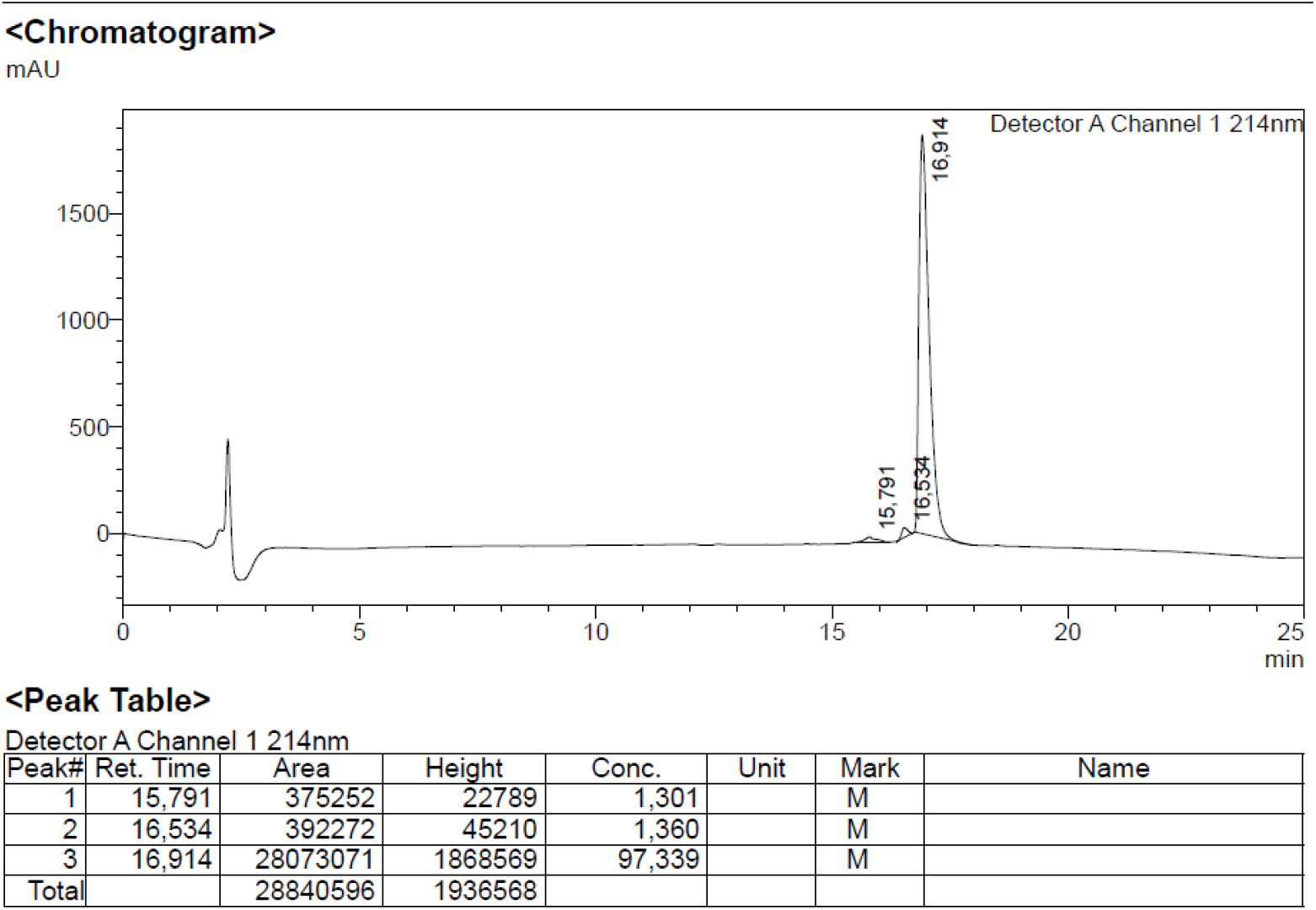
Analytical HPLC of **MF6**

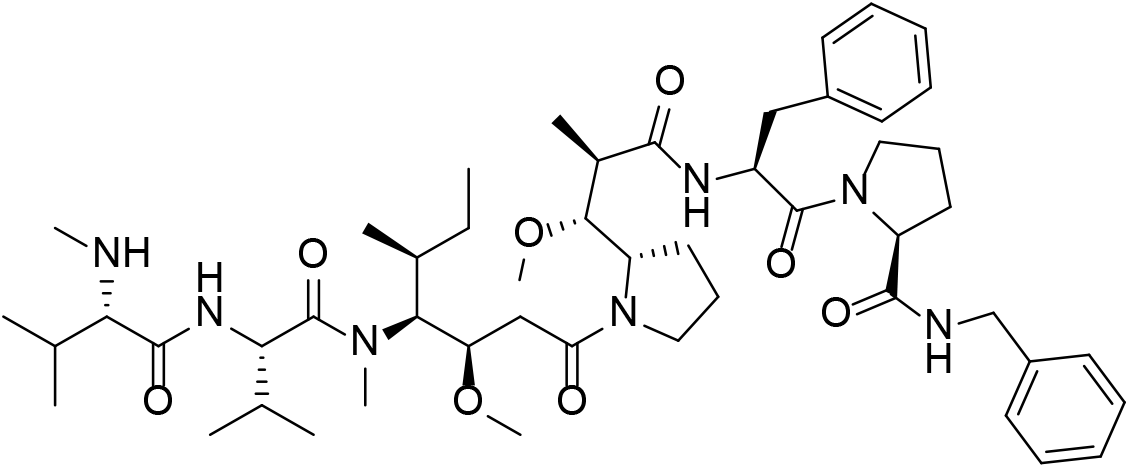

##### MF7

To a solution of Boc-*L*-Pro-OH (10.00 mg, 0.05 mmol, 1 eq.) in DCM, CDI (9.040 mg, 0.056 mmol, 1.2 eq.), Oxyma Pure (13.210 mg, 0.093 mmol, 2 eq.) was added and stirred at room temperature for 5 min. Benzylamine (5.194 mg, 5.04 μL, 0.056 mmol, 1.2 eq.) was then added dropwise. The mixture was stirred at room temperature for 4 h. After fully conversion of Boc-*L*-Pro-OH, the reaction mixture was added to DCM (5 mL) in separatory funnel. The organic layer was washed with 5 mL of NaHCO_3_(sat.) then water (5 mL) and 5 mL of 0.5 % HCl. Organic layer was evaporated to obtain crude product. Dioxane (1 mL) was added to the crude product, followed by 4N HCl in dioxane (1 ml) and resulting was stirred at room temperature for 4h. The reaction mixture was then concentrated and dried for 24h to afford crude free amine.

**Fmoc-MMAF** (6 mg, 0.0062 mmol, 1 eq.), HATU (9.5 mg, 0.025 mmol, 4 eq.), Oxyma Pure (1.8 mg, 0.0124 mmol, 4 eq.) was dissolved in 1 ml of 20 % of collidine. The mixture was then added to a solution of the above-mentioned crude free amine (pre-dissolved in 1 ml 20% collidine in DMF) and stirred for 24h. After consumption of **Fmoc-MMAF** (TLC), the reaction mixture was added to 10 ml of ethyl acetate and washed with 10 mL of 5 % NaHCO_3_ and 10 mL sat. oxalic acid. The organic layer was concentrated to afford crude **Fmoc-MF7**. The crude was washed with cold ethyl ether (3 times), pentane (3 times) and used directly for the final deprotection.

To a solution of crude **Fmoc-MF7** in 0.5 ml DMF, 1ml of 20% piperidine in DMF was added and stirred for 5 min. The reaction mixture was concentrated under reduced pressure to afford crude **MF7**. The crude was washed with cold ethyl ether (3 times), pentane (3 times) and purified by preparative-HPLC to afford 2.2 mg of final product. HRMS of **MF7** Cal. For C_51_H_80_N_7_O_8_ [M+H]^+^ 918.6068, found 918.6066.

**Figure S18.**
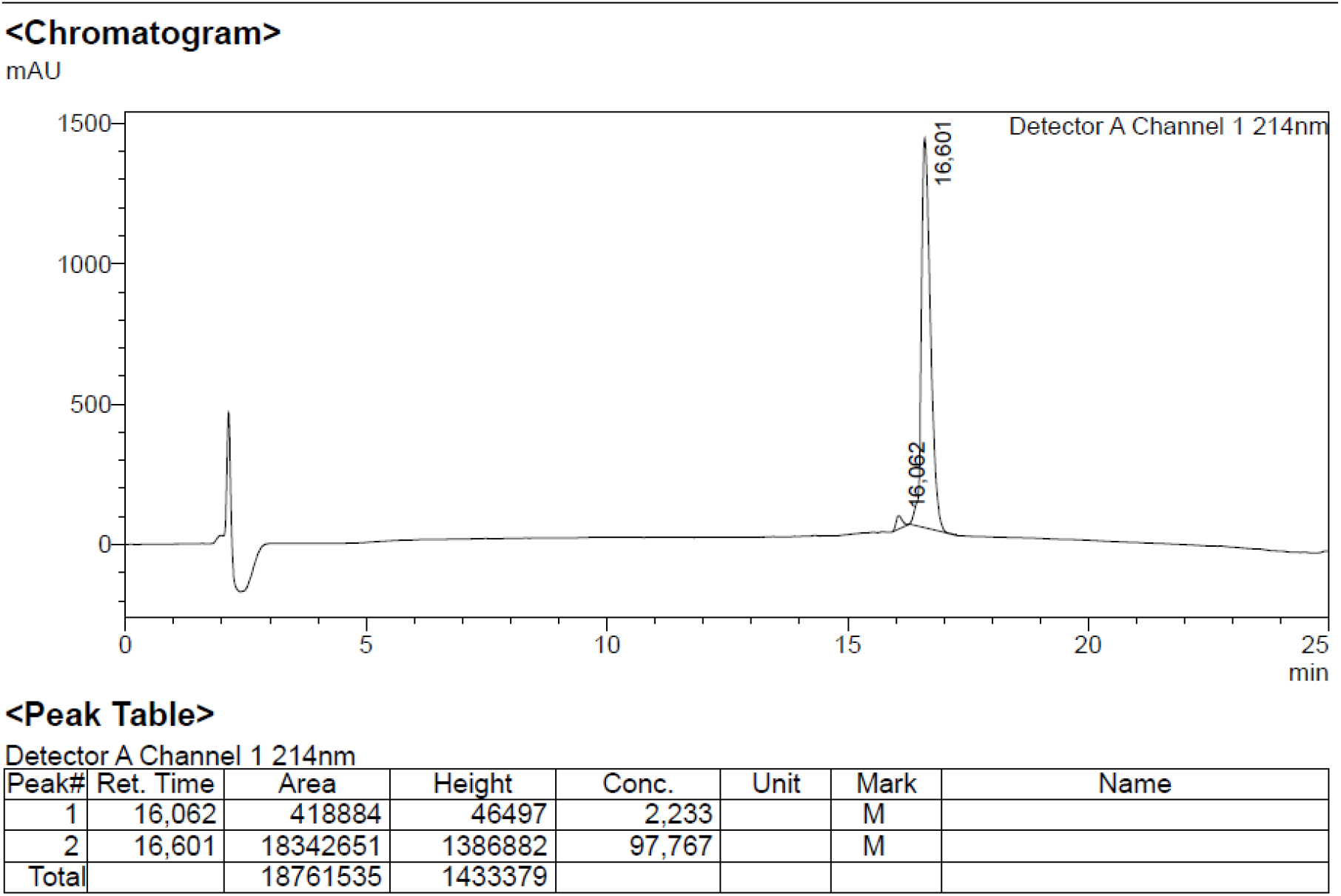
Analytical HPLC of **MF7**

#### Solid-phase peptide synthesis MFR1-6

##### General procedure for synthesis of MFR1-6 library compounds

Peptides **MFR1-6** were synthesized through solid phase peptide synthesis using a 2-chlorotrityl (2CT) resin and standard Fmoc protecting group strategy. 2CT resin (1.6 mmol/g, 31 mg, 0.0496 mmol, 1 equiv.) was placed in a syringe. The resin was washed thrice with DMF and thrice with DCM. A solution of First Fmoc-amino acid (0.0622 mmol, 1.3 equiv.) and DIPEA (11 μL, 0.0643 mmol, 1.3 equiv.) in DCM (1 mL) was added to the resin and stirred for 5 min. Another aliquot of DIPEA (30 μL, 0.175 mmol, 3.5 equiv.) was added to the reaction, which was stirred for 1 hour at room temperature. Methanol (50 μL) was added to the reaction, which was then stirred for 30 min. The resin was then washed thrice with DMF and thrice with DCM.

The following couplings followed the same pattern. First Fmoc removal of resin by reaction with piperidine/DMF (1:4) for 3 min, 10 min and 10 min. Then a solution of the next amino acid (0.133 mmol, 2.7 equiv.) was activated in DMF with oxyma (0.0915 mmol, 1.8 equiv.) and DIPC (13 μL, 0.0834 mmol, 1.7 equiv.) for 5 min with stirring, upon which it was added to the resin. The resin was stirred at room temperature for 1.5 h and washed thrice with DMF and thrice with DCM. After each coupling, a Kaiser test was performed to confirm coupling to all free amines before deprotection and subsequent coupling.

The last coupling was performed with Fmoc-MMAF (1) (52 mg, 0.0545 mmol, 1.1 equiv.) which was pre-activated with oxyma (7 mg, 0.0493 mmol, 1 equiv.) and DIPC (7 μL, 0.0496 mmol, 1 equiv.) in DMF for 5 min. The solution was added to the resin and stirred overnight. The resin was nevertheless washed with DMF thrice and DCM thrice. Fmoc was removed by reaction with piperidine/DMF (1:4) for 3 min, 10 min, and 10 min. The peptide was cleaved from the resin with a 2 mL TFA/H2O/DTT (90:5:5) for 1 hour followed by concentration and trituration. The crude was purified by preparative-HPLC yielding the pure product.

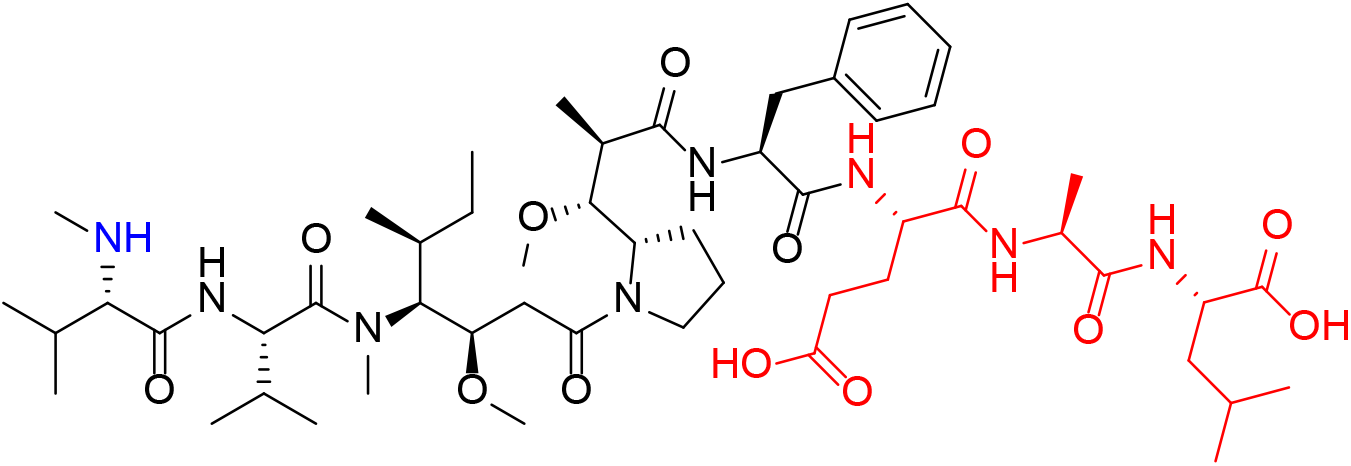

##### MFR1

General procedure, amino acids used: Glu, Ala, Leu. 5.9 mg after preparative-HPLC

**Figure S20.**
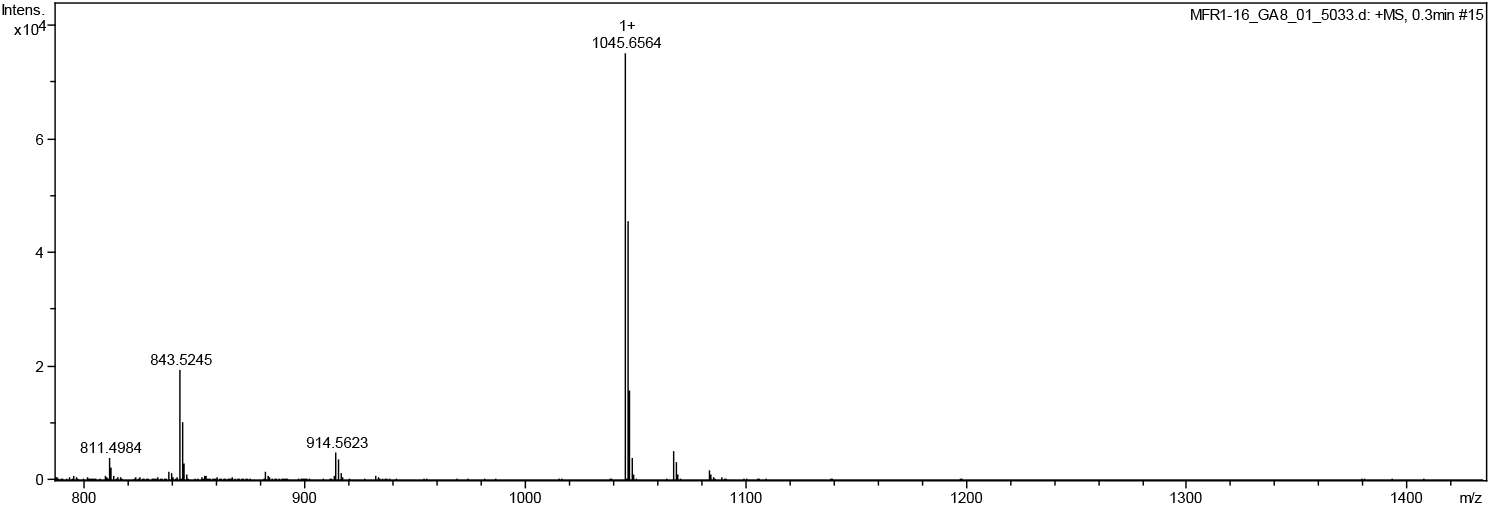
HRMS of **MFR1**, Cal. For C_53_H_89_N_8_O_13_ [M+H] ^+^ 1045.6549, found1045.6564

**Figure S21.**
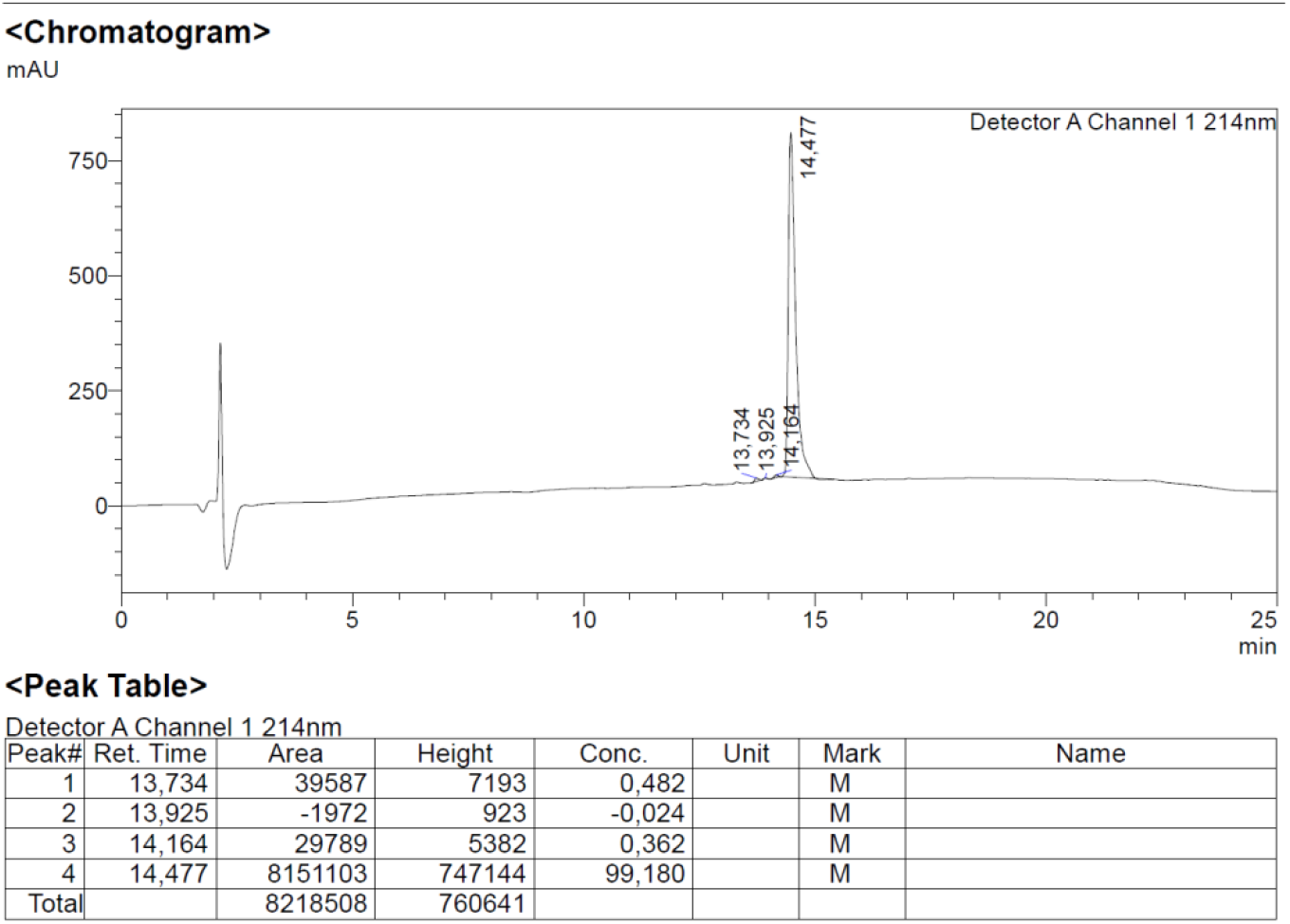
Analytical HPLC of **MFR1**

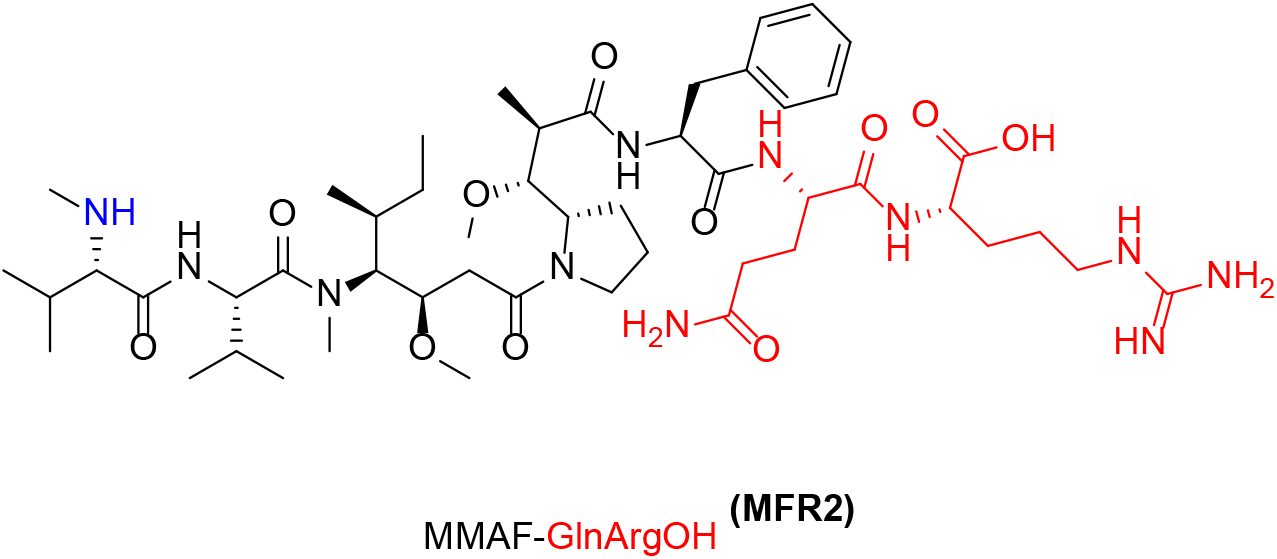

##### MFR2

General procedure, amino acids used: Gln, Arg. 2.8 mg after preparative-HPLC.

**Figure S22.**
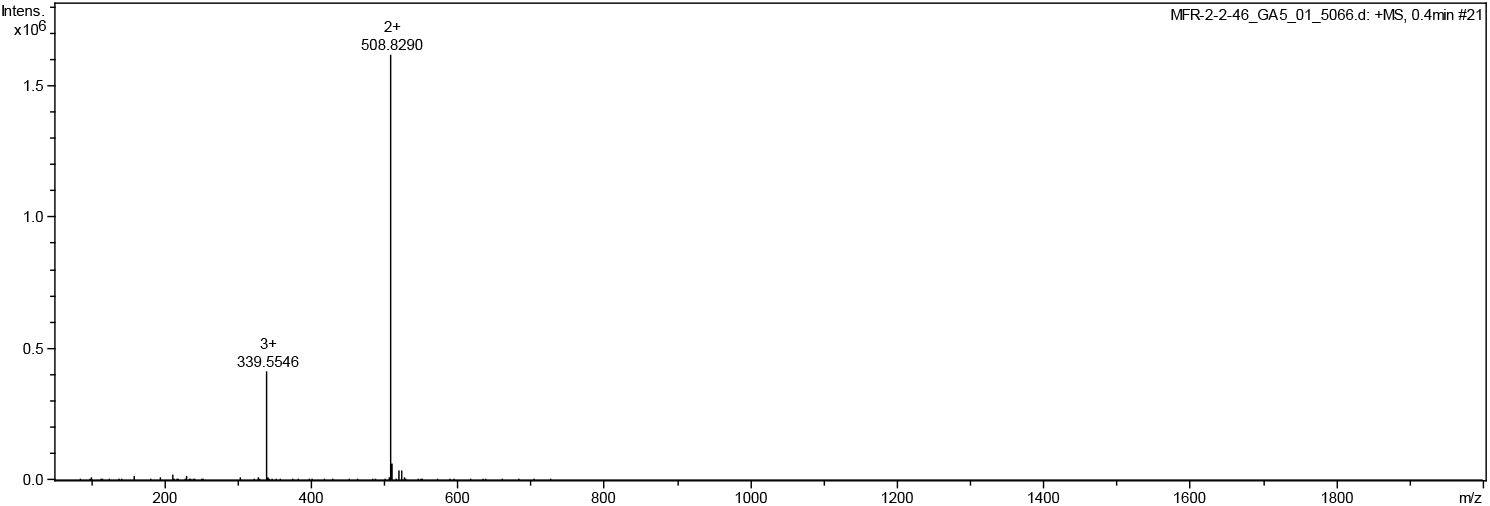
HRMS of **MFR2** Cal. For C_50_H_86_N_11_O_11_ [M+1] ^+^ 1016.6508, found 508.8290 [M/2]^2+^.

**Figure S23.**
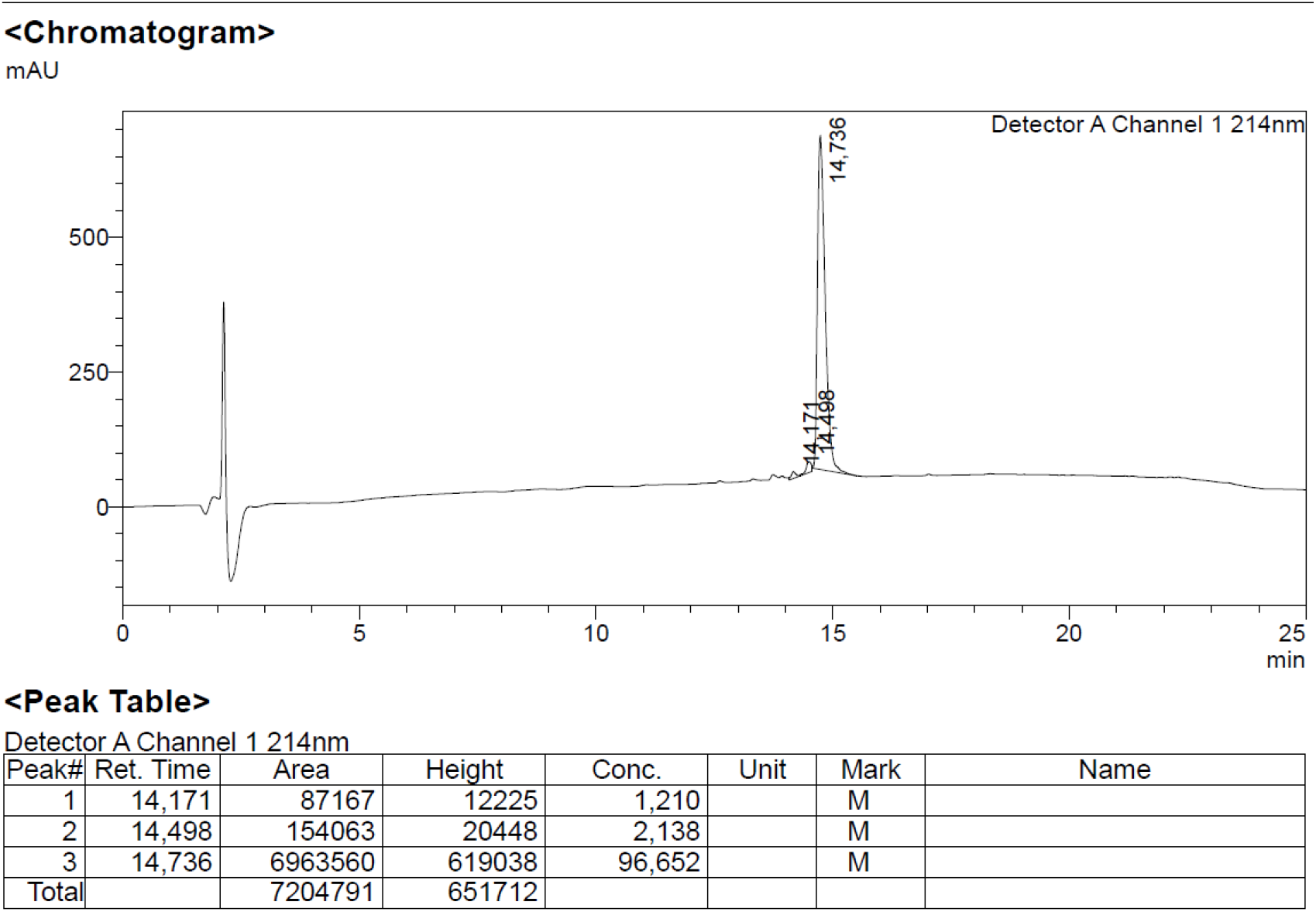
Analytical HPLC of **MFR2**

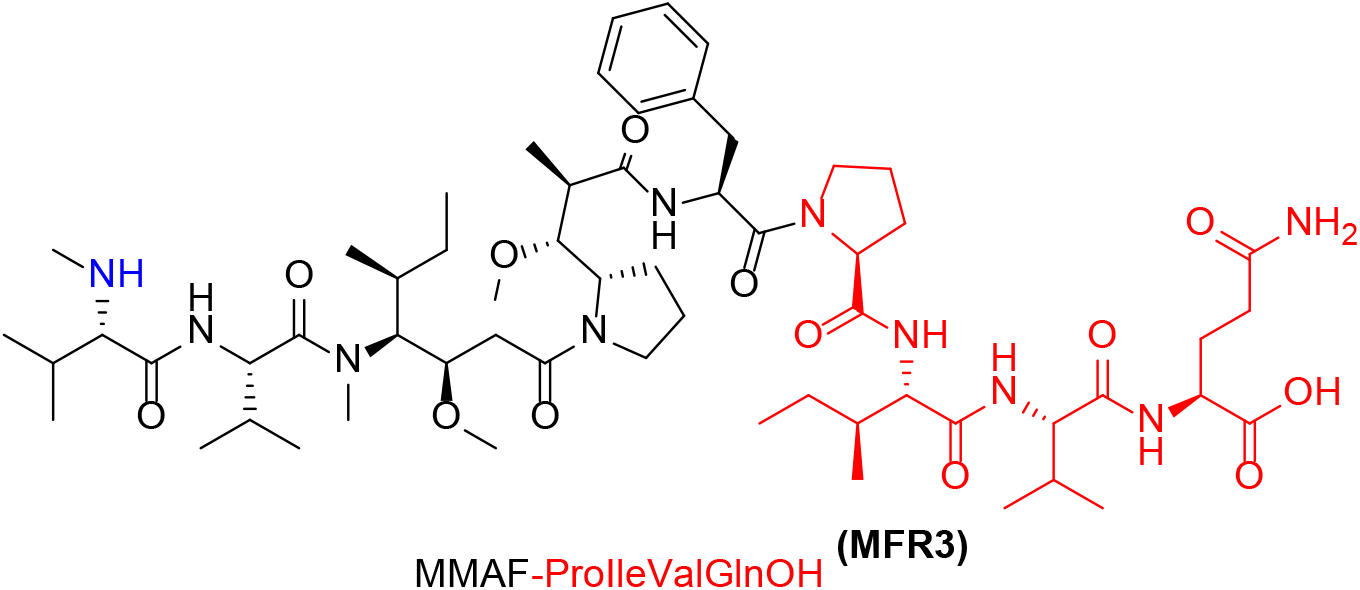

##### MFR3

General procedure, amino acids used: Pro, Ile, Val, Gln. 5.8 mg after preparative-HPLC

**Figure S24.**
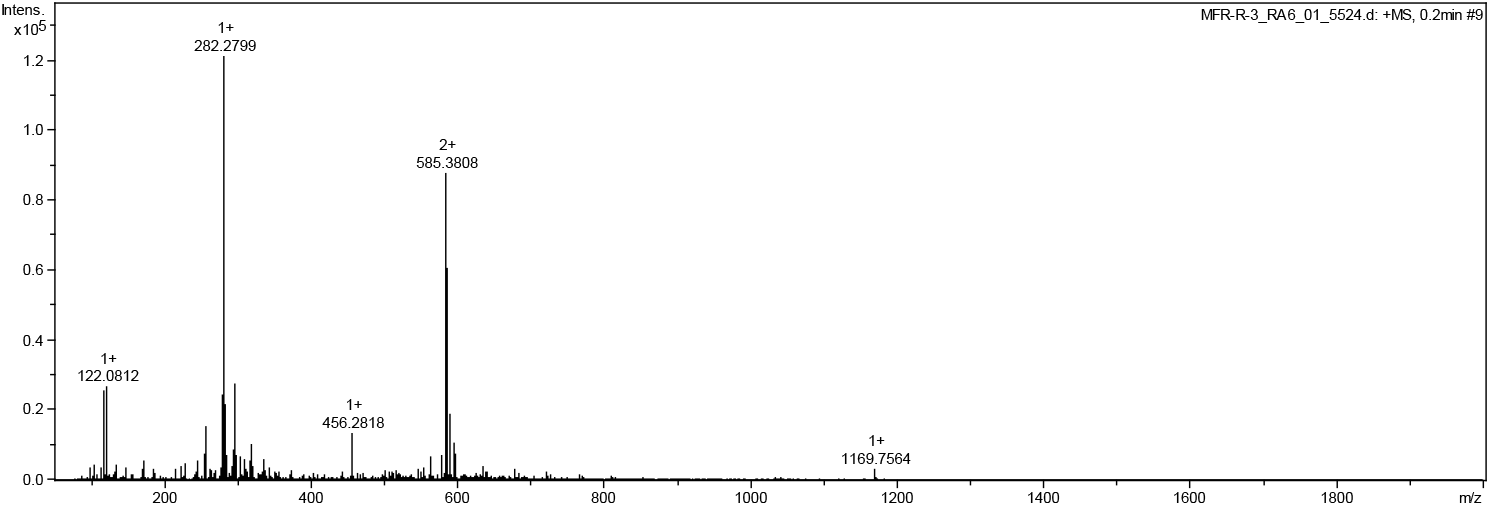
HRMS of **MFR3** Cal. For C_60_H_101_N_10_O_13_ [M+H] ^+^ 1169.7550, found 1169.7564

**Figure S25.**
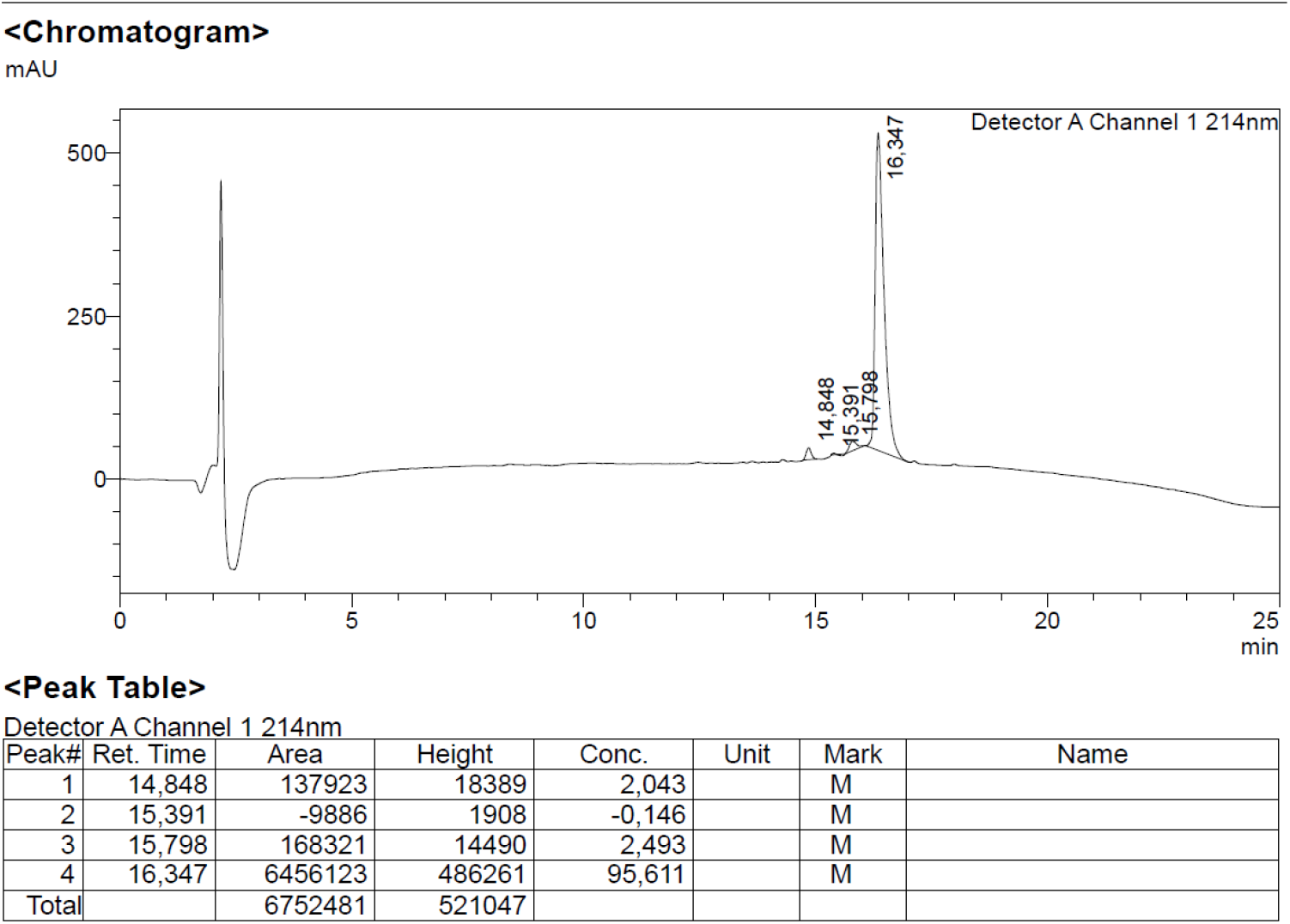
Analytical HPLC of **MFR3**

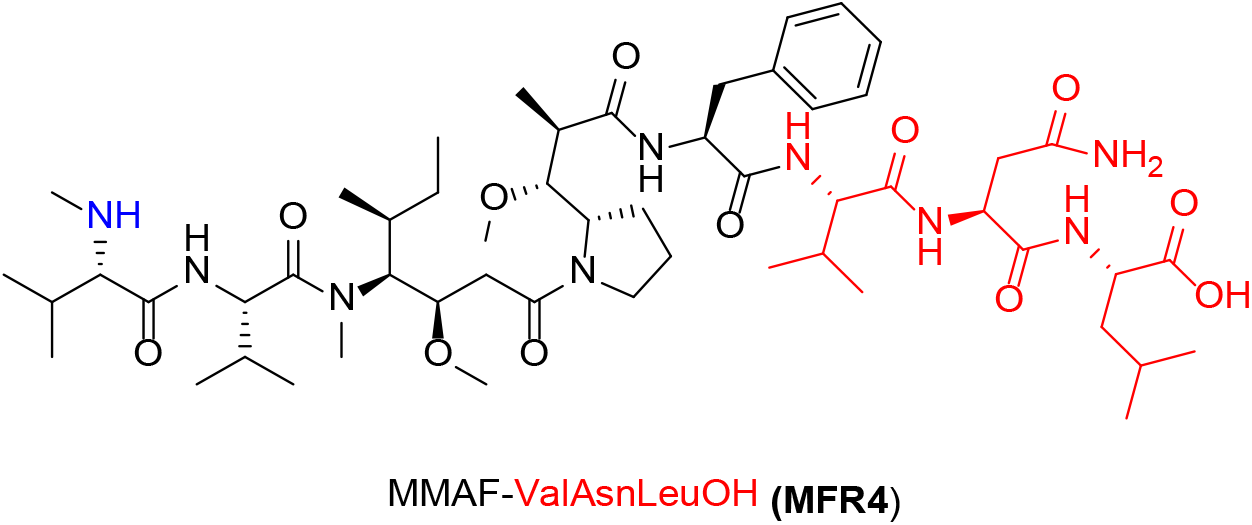

##### MFR4

General procedure, amino acids used: Val, Asn, Leu. 2.7 mg after preparative-HPLC

**Figure S26.**
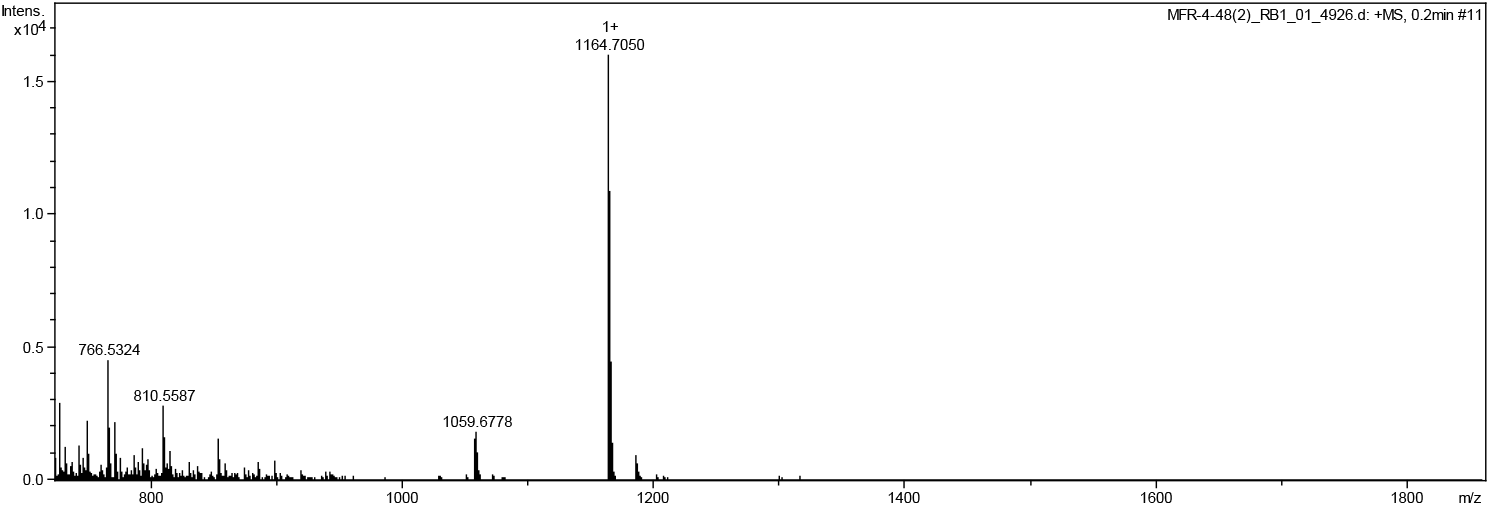
HRMS of **MFR4** Cal. For C_54_H_92_N_9_O_12_ [M+H]^+^ 1059.6937, found 1059.6778.

**Figure S27.**
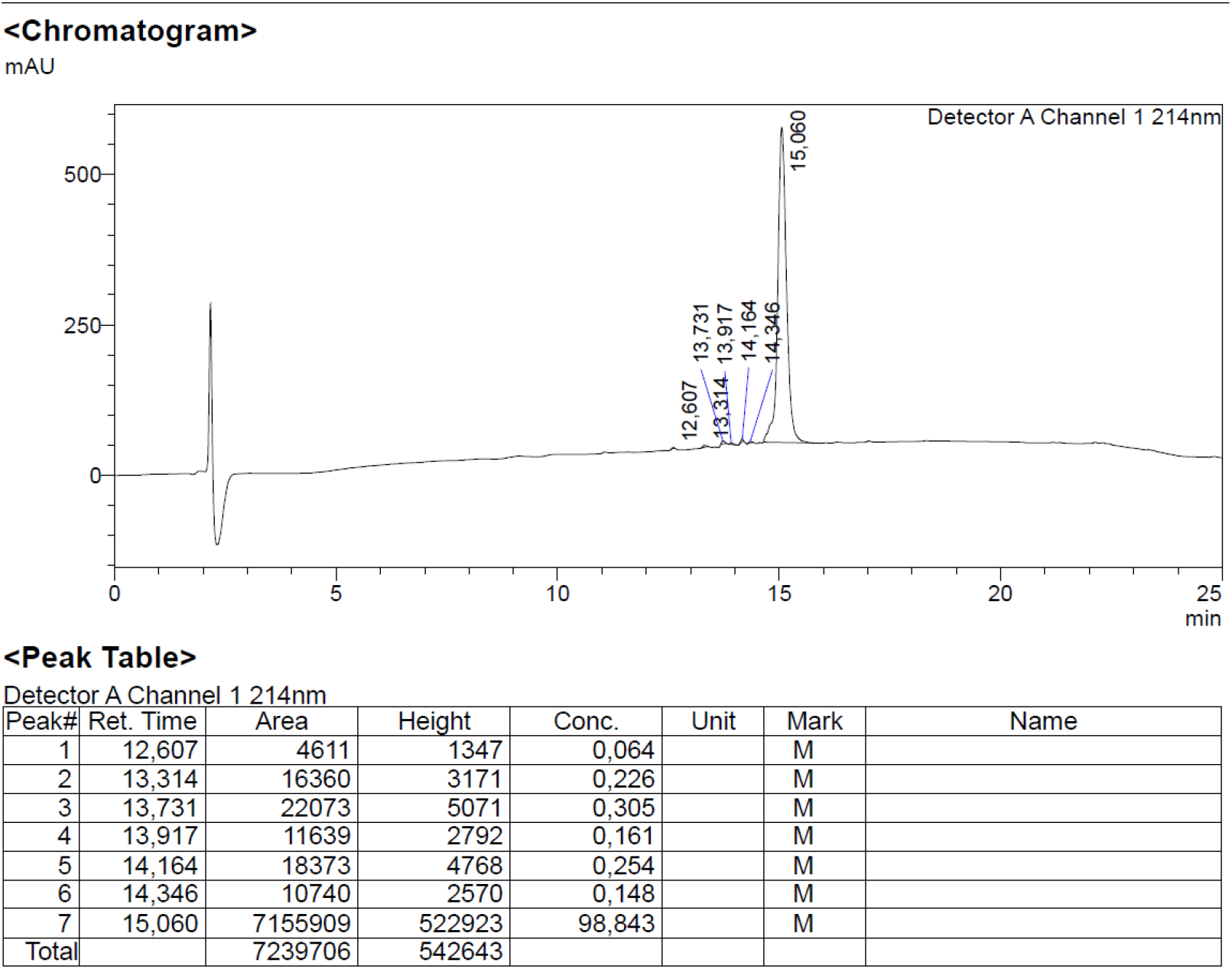
Analytical HPLC of **MFR4**

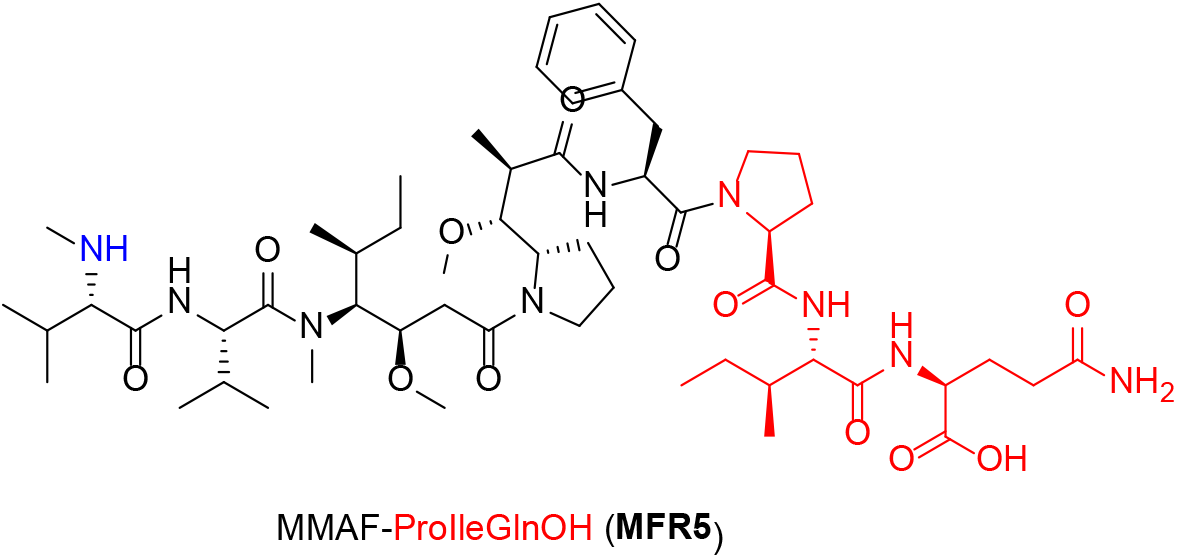

##### MFR5

General procedure, amino acids used: Pro, Ileu, Gln. 6.2 mg after preparative-HPLC

**Figure S28.**
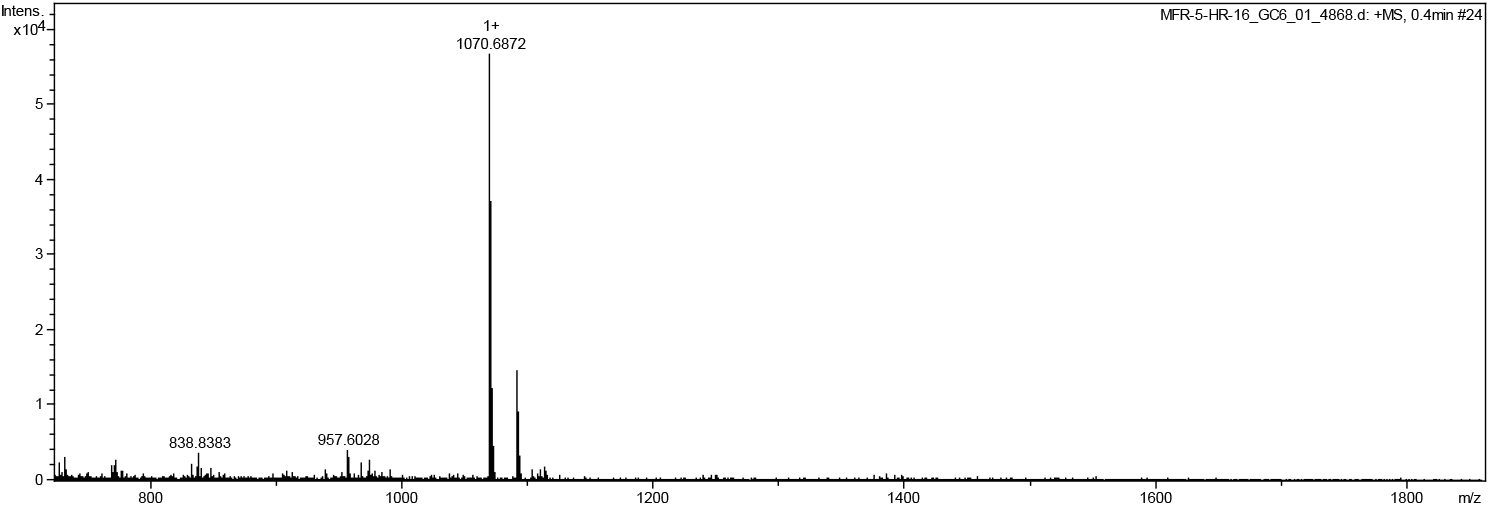
HRMS of **MFR5** Cal. For C_55_H_92_N_9_O_1_2 [M+H]^+^ 1070.6865, found 1070.6872

**Figure S29.**
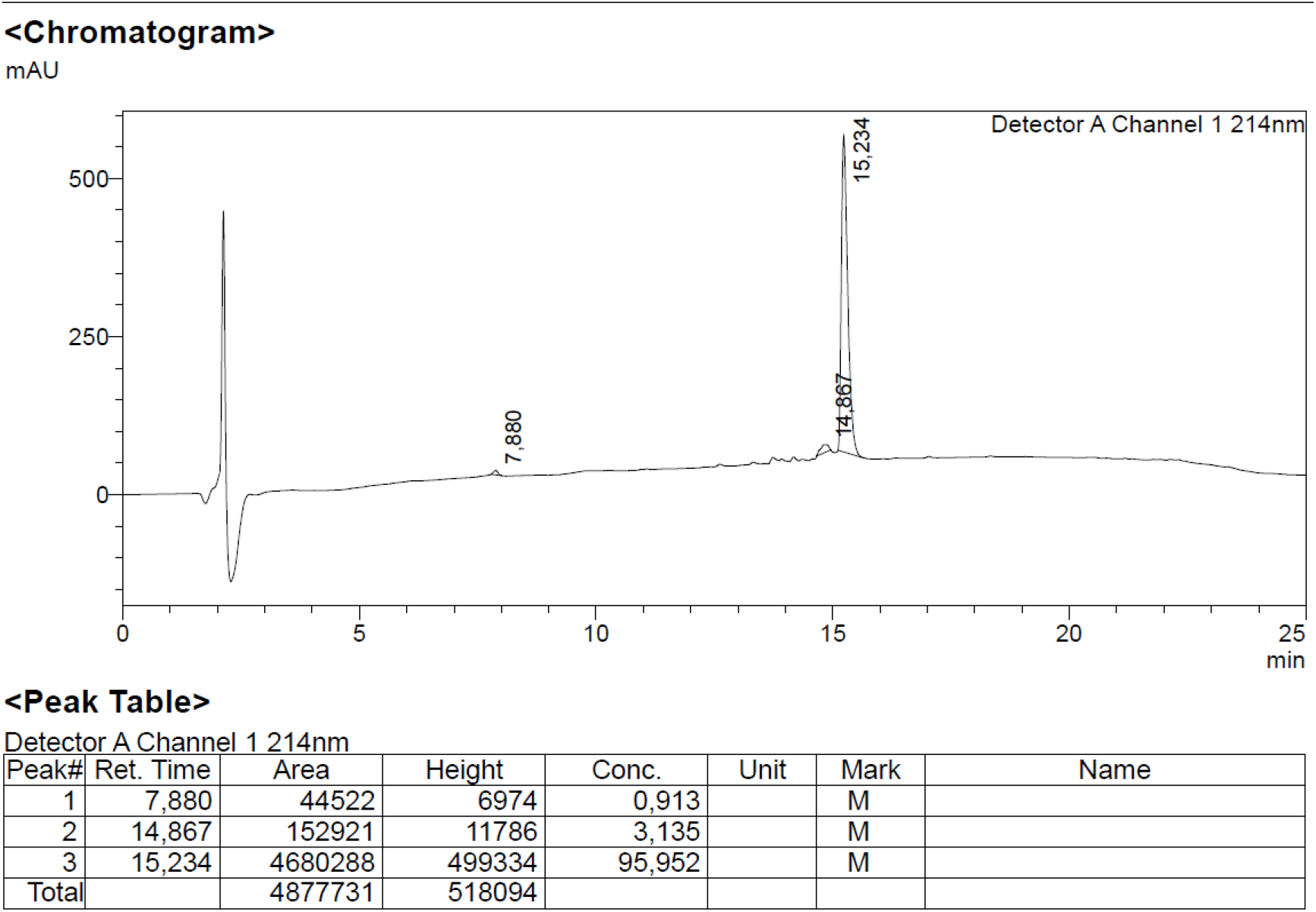
Analytical HPLC of **MFR5**

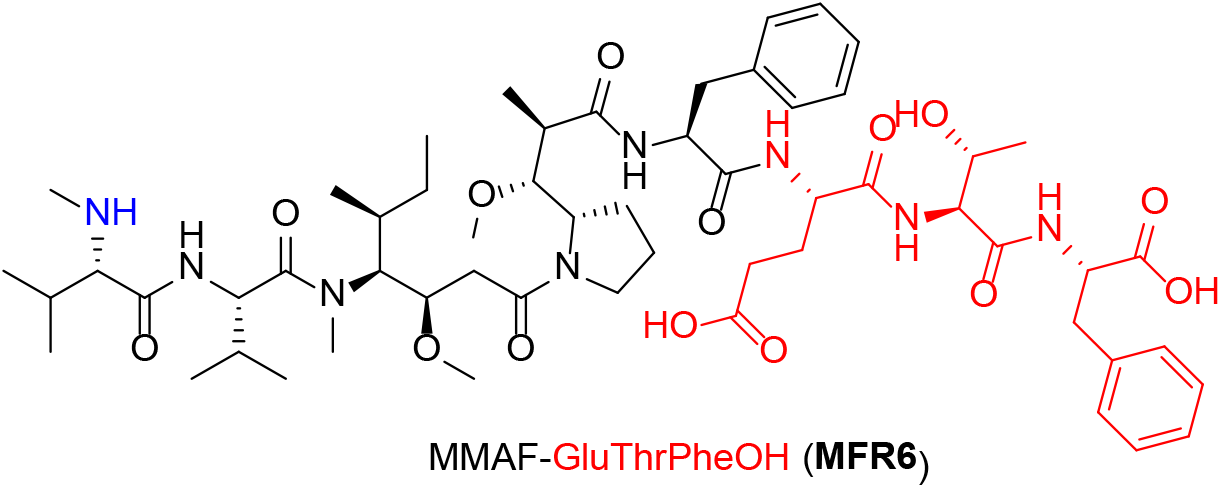

##### MFR6

General procedure, amino acids used: Glu, Thr, Phe. 4.8 mg after preparative-HPLC

**Figure S30.**
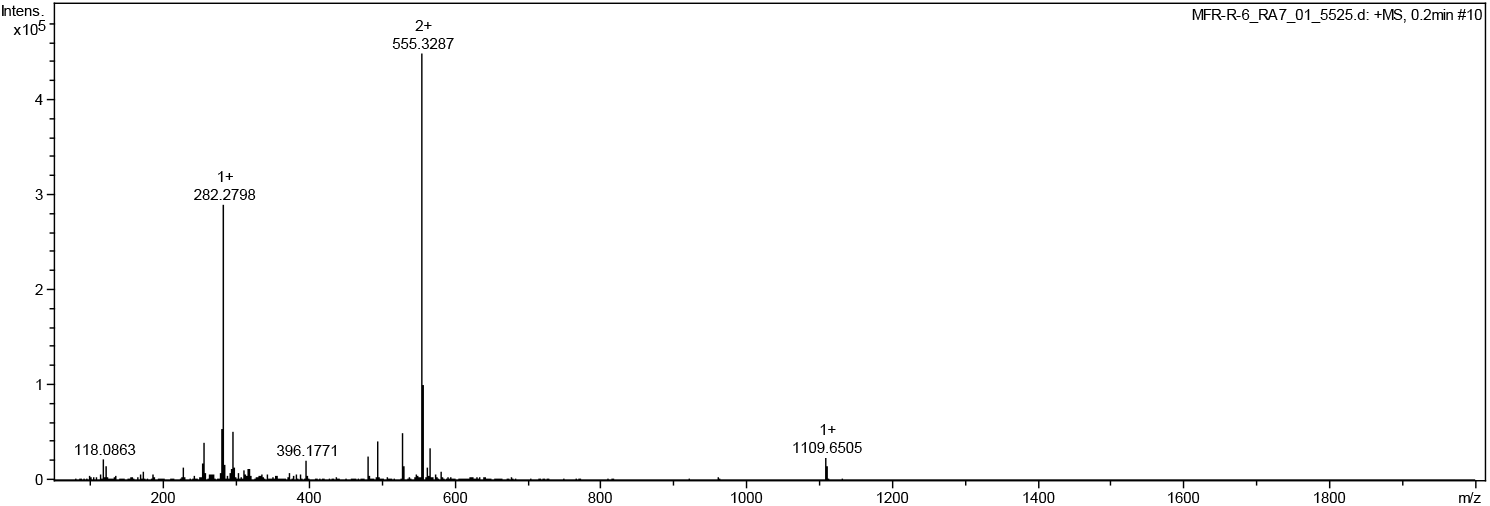
HRMS of **MFR6** Cal. For C_57_H_89_N_8_O_14_ [M+H]^+^ 1109.6498, found 1109.6505

**Figure S31.**
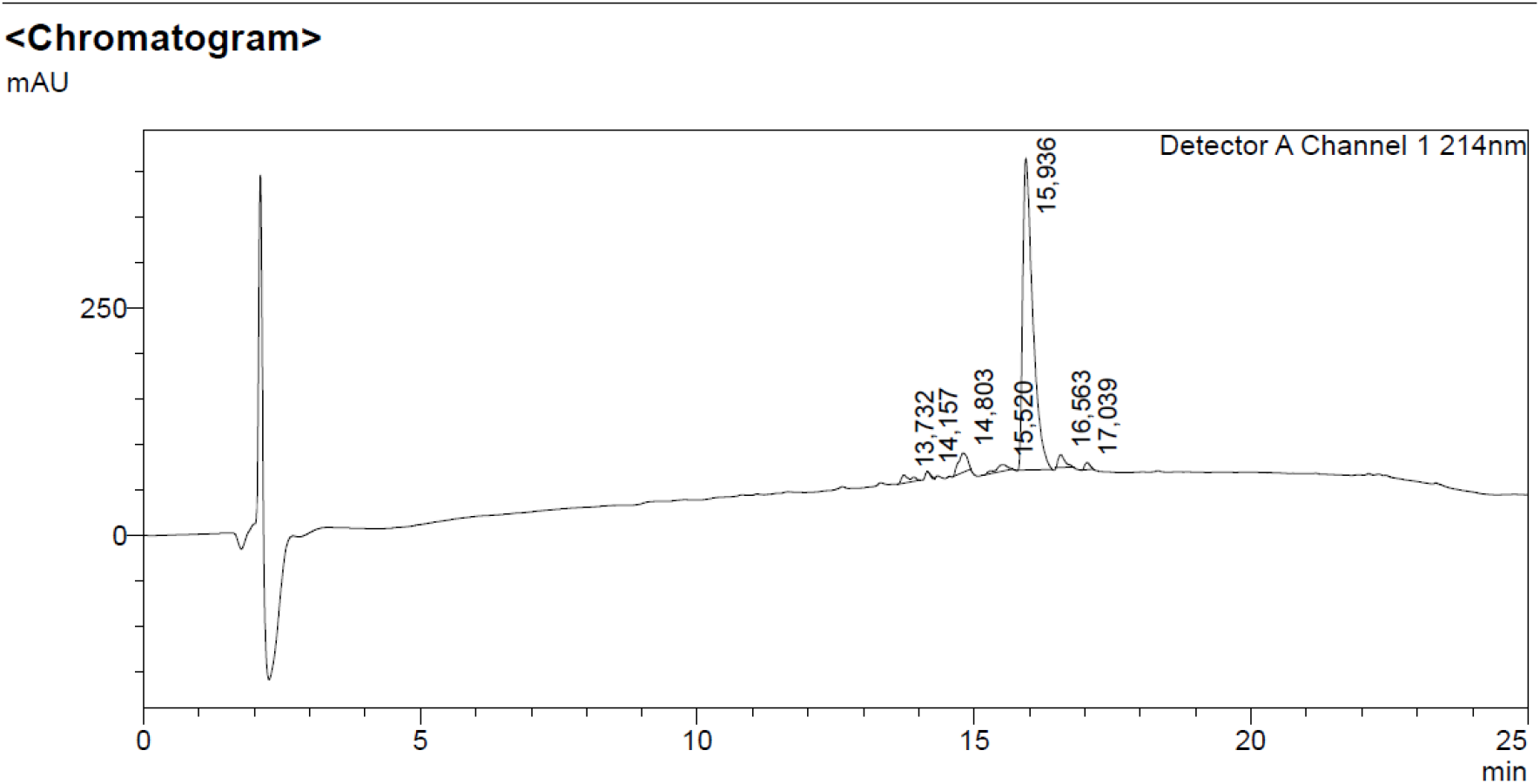
Analytical HPLC of **MFR6**

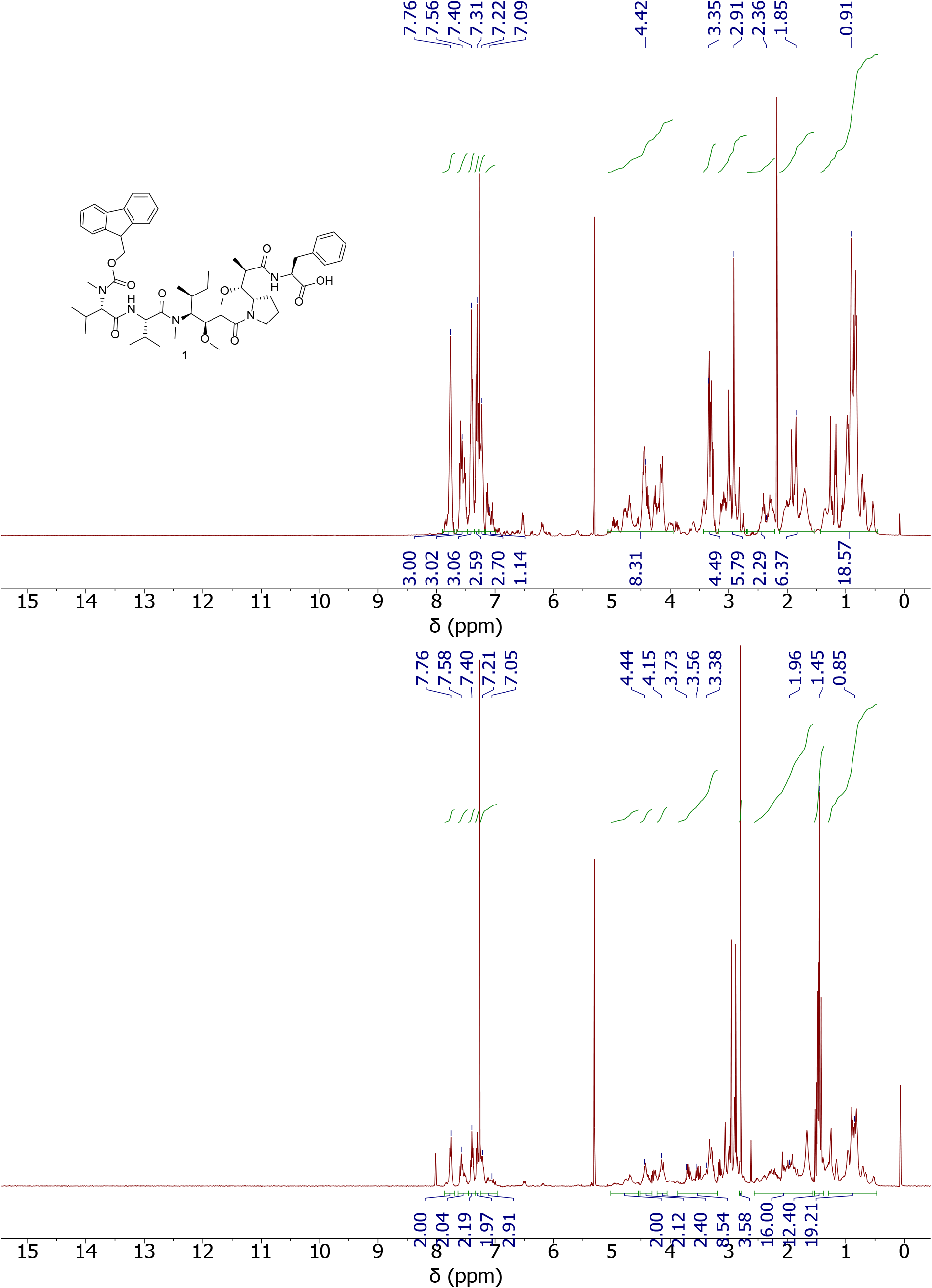

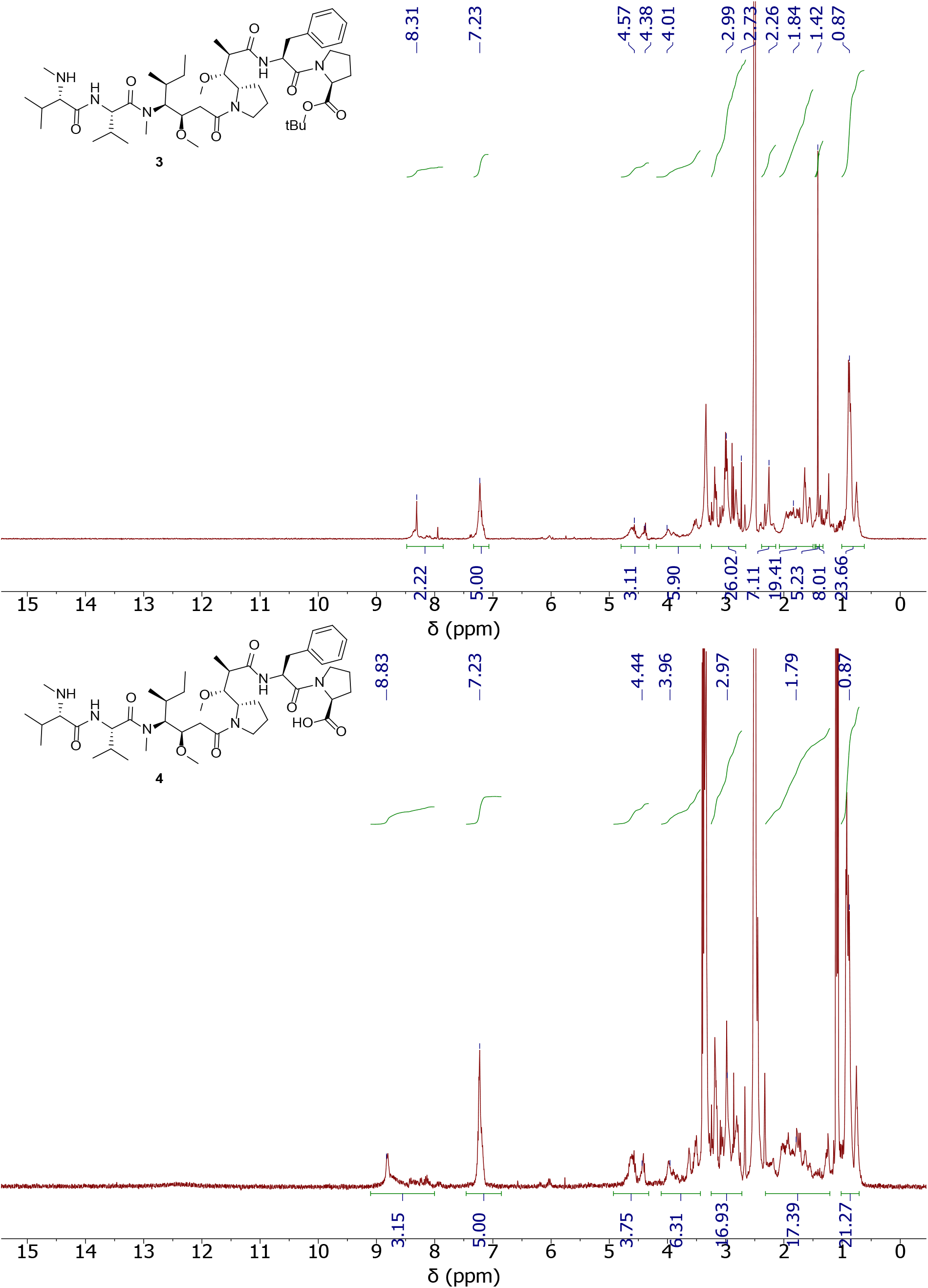

